# The germline-specific region of the sea lamprey genome plays a key role in spermatogenesis

**DOI:** 10.1101/2021.09.24.461754

**Authors:** Tamanna Yasmin, Phil Grayson, Margaret F. Docker, Sara V. Good

## Abstract

The sea lamprey genome undergoes programmed genome rearrangement (PGR) in which ∼20% is jettisoned from somatic cells soon after fertilization. Although the role of PGR in embryonic development has been studied, the role of the germline-specific region (GSR) in gonad development is unknown. We analysed RNA-sequence data from 28 sea lamprey gonads sampled across life-history stages, generated a genome-guided *de novo* superTransciptome with annotations, and identified genes in the GSR. We found that the 638 genes in the GSR are enriched for reproductive processes, exhibit 36x greater odds of being expressed in testes than ovaries, show little evidence of conserved synteny with other chordates, and most have putative paralogues in the GSR and/or somatic genomes. Further, several of these genes play known roles in sex determination and differentiation in other vertebrates. We conclude that the GSR of sea lamprey plays an important role in testicular differentiation and potentially sex determination.

## Introduction

The genetic structure and composition of germline and somatic cells typically remain constant throughout an organism’s life span. However, under some conditions (e.g., cancer) or in some taxa, the genetic composition of cells varies by type and/or developmental stage^1,2^. Included in this is the unusual process of programmed genome rearrangement (PGR), in which either portions of chromosomes (chromosomal diminution) or entire chromosomes (chromosomal elimination) are removed during embryonic development, thereby reducing the genomic content of descendent cells by up to 90%^3^. Although the frequency of PGR across metazoans is unknown, it has been observed in more than 100 vertebrate and invertebrate species from nine major taxonomic groups^3^, including in lampreys^4–7^. In sea lamprey (*Petromyzon marinus*), flow cytometric measurements of DNA content in the germline (testes) vs. somatic (blood) cells indicate that ∼20% (∼500 Mb) of the germline genome is eliminated during PGR^8^. Further studies have shown that PGR in sea lamprey, which occurs ∼3 days post-fertilization (dpf), shares conserved features with PGR in other agnathan lineages^4–7^. This event involves chromosomal elimination of repetitive and single-copy sequences and is enriched for genes involved in development or germline maintenance^6,9^. However, further research on the possible function of the germline-specific regions (GSR) in gonad development is needed.

Many hypotheses have been posited regarding the biological significance of PGR, including gene silencing, dosage compensation, position effects on gene expression, germline development, and sex determination^1,10–13^. In sea lamprey, it has been suggested that PGR permits the expression of genes beneficial to the germline during the early stages of embryonic development^6,8^, consistent with the high levels of gene silencing observed for genes in the GSR^3,9^. In the zebra finch (*Taeniopygia guttata*), chromosomal diminution of a germline-restricted chromosome (GRC) occurs during early embryonic development; the genes in the GRC have higher expression in the ovary than the testis, and the GRC is later eliminated from mature sperm, being transmitted only through the oocytes^14,15,16^.

In sexually reproducing taxa without PGR, primordial germ cells (PGCs) are formed early during embryogenesis. PGCs typically develop through the coordination of three developmental cues: suppression of ongoing somatic differentiation, repression of DNA methylation, and inhibition of cell proliferation^17^. Once defined, PGCs subsequently exhibit tightly coordinated gene expression that leads to germ cell development and differentiation in both sexes. In lampreys, however, the germline cells are specified at fertilization, and somatic cell delineation occurs afterward, ∼3 dpf, when PGR is initiated^5^. This intriguing reversal of events is heightened by the ongoing enigma of their sex determination. Lampreys do not have heteromorphic sex chromosomes and there is no evidence to date of genomic differences between males and females; sex may be determined by genetic factors in the germline genome, environmental factors, or a combination of the two (reviewed by^18^).

Here, we used RNA-sequence (RNA-seq) data from 28 sea lamprey gonads sampled at different life-history stages and in both sexes to generate a gonadal superTranscriptome and examined the function, expression, and evolutionary relationships of sex-biased genes, particularly in the GSR. We identified 638 germline-specific genes (GSGs), many of which were present in multiple germline-specific paralogues pertaining to 163 unique gene names that were, overall, very highly expressed during spermatogenesis, but lowly expressed during oogenesis and in undifferentiated larvae. The observation that the genes in the GSR appear to be present in undifferentiated larvae and females but are expressed at low levels suggests that the male-specific expression is due to regulatory changes, as opposed to there being a male-specific germline sequence. Further, we found that ∼55% of the GSGs also have paralogous copies in the somatic genome and ∼19% have putative orthologues in other taxa including, most importantly, a core set of conserved genes involved in sex determination and spermatogenesis. Using publicly available RNA-seq data from 1–5 dpf embryos, we found that the genes expressed during gonadogenesis are either not expressed or lowly expressed during early embryo formation. Collectively, these results suggest that a major role of the GSR is in testicular differentiation and probably sex determination. PGR in sea lamprey may serve to reduce conflict of genes under sexual selection, a hypothesis further supported by the highly duplicated nature of genes in the GSR and their association in sexual differentiation and determination pathways in other taxa (see^3^).

## Results and discussion

### GSGs show predominantly male-biased expression and have a key role in gametogenesis

We used RNA-seq data from 28 sea lamprey gonads sampled across a range of developmental stages to generate a gonadal superTranscriptome using the Necklace pipeline^19^. Stages included undifferentiated larvae, female larvae following the onset of oogenesis and sexually mature (adult) females, prospective male larvae (i.e., those in which the gonad was still histologically undifferentiated but which were beyond the size at which ovarian differentiation is complete), males undergoing testicular differentiation following the onset of metamorphosis, and sexually mature (adult) males (see Supplementary Fig. 1 and Supplementary Table 1). This revealed a large number of genes that were highly expressed during male but not female gonad development; these genes were physically linked and mapped to chromosome 81 and many unplaced scaffolds based on the Vertebrate Genome Project (VGP) reference assembly. Thus, we sought to define which of the genes in our gonadal superTranscriptome mapped to the GSR.

The GSR in sea lamprey was identified for an earlier release of the sea lamprey germline assembly (www.stowers.org)^20^. Thus, we used a modified version of the DifCover pipeline used for that analysis^21^ to define the coordinates of GSR in the VGP assembly. Accordingly, GSRs were designated as regions in which the read coverage of sperm DNA was > 2-fold more than the read coverage of blood DNA. Based on the VGP reference genome, a total of 5253 genomic intervals were mapped by DifCover, of which 919 segments had an enrichment score (log2(standardized sperm coverage/blood coverage) greater than 2. The total span of the GSR-inferred regions consisted of more than 27 Mbps (Supplementary Table 2, Supplementary Fig. 2).

We then used the segment enrichment scores to assign genes from our gonadal superTranscriptome to either the GSR or somatic genomes. Using an earlier scaffold-based assembly of the sea lamprey germline genome (available at SIMRbase), Smith et al., (2018) identified ∼13Mbps including 356 protein-coding genes in the GSR^20^. On the other hand, using the VGP assembly which consists of 85 chromosomes and 1195 unassembled scaffolds, we assigned the entirety of chromosome 81 as well as 177 scaffolds (Supplementary Fig. 3) to the GSR, while the remaining 84 chromosomes and 1018 scaffolds were not germline-enriched, suggesting that they are found in the somatic genome. In total, 638 genes from our gonadal superTranscriptome mapped to the GSR; these 638 genes corresponded to only 163 unique gene names based on our combined Trinotate and reference genome annotation pipeline (Supplementary Table 3), with approximately half of the GSGs occurring in a single copy but the other half occurring in 2–77 duplicated copies (Supplementary Table 4, Supplementary Fig. 4). Importantly, however, none of the GSR enriched scaffolds or chromosomes showed overlap between the GSR and the somatic regions. This supports previous work that determined that PGR in lampreys is more likely to involve chromosome elimination than diminution^9^.

The expression analysis of the GSGs revealed that out of 638 GSGs, 409 genes (64% of the genes in the GSR) are moderately to highly expressed in one or more stages of the developing testis, but only a few GSGs are expressed in undifferentiated larval gonads (Fig. 1a, see Supplementary Fig. 5 for full heatmap). Functional enrichment analysis of the GSGs from the gonadal superTranscriptome indicate that they are involved in 26 pathways of which *wnt* signaling and *E-Cadherin* signaling pathways each represented 16.4% of the total hits (Supplementary Fig. 6). Other critical pathways include the insulin/insulin growth factor (*igf*) pathway, gonadotropin releasing hormone receptor (*gnrhr*) pathway, transforming growth factor beta (*tgfb*) signaling pathway, and fibroblast growth factors (*fgf)* pathway, which contained 2.7%, 2.7%, 2.7%, and 1.4% of all hits, respectively (Supplementary Fig. 6). Next, we analyzed the GO terms associated with genes in the GSR to obtain further insight into their molecular function (Supplementary Table 5). Using an overrepresentation test, we found that the highest FDR terms were associated with reproductive system development, positive and negative regulation of cell population proliferation, ovarian follicle development, oogenesis and spermatogenesis. Collectively, this demonstrates that the functional ontology of GSGs is enrichment for GO terms related to reproductive developmental processes (Fig. 1b, Supplementary Table 6).

**Fig. 1:**
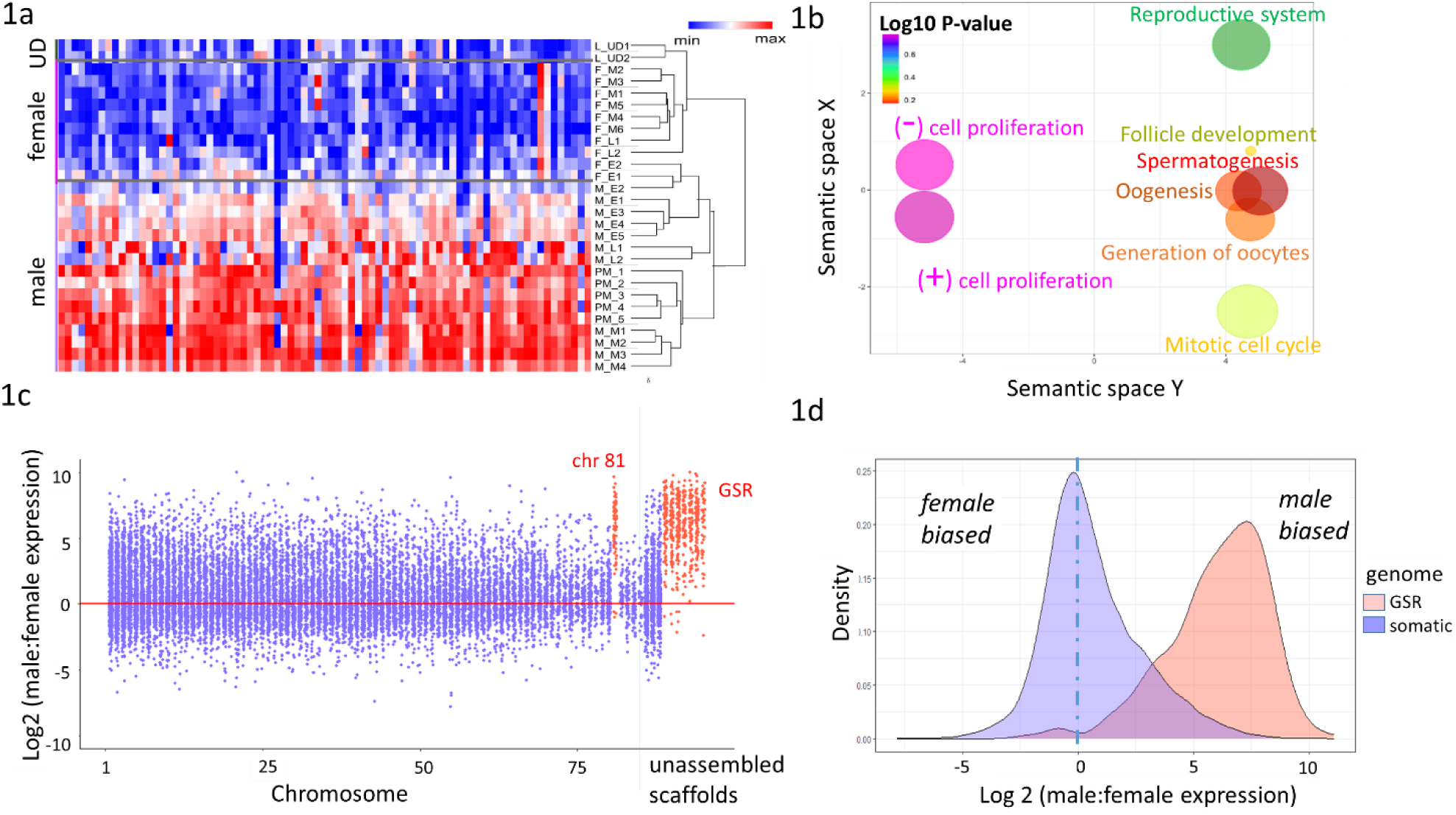
Identification of genomic location of the germline-specific genes (GSGs) in the VGP genome and their expression in the sea lamprey gonadal samples used in this study. **a)** Heatmap showing GSG expression pattern of all samples used in the study; UD stands for undifferentiated larvae (see Supplementary Fig. 3 for full heatmap) **b)** Gene ontology term enrichment analysis of the GSGs where colours indicate the log10 of the false discovery rate-corrected *P*-value (PANTHER overrepresentation test, with a Fisher exact test for significance and filtering using a false discovery rate of 0.05); circle size denotes fold enrichment above expected values. **c)** Scatterplot showing the log_2_(male:female) normalized gene expression across all chromosomes and concatenated scaffolds in the VGP assembly of the sea lamprey genome; regions identified as belonging to the GSR are coloured in red, while those in the somatic genome are coloured in blue. **d)** Density plot of the log2(male:female) ratio of normalized gene expression. When x = 0, average expression across all females = males. For genes in the somatic genome, the density peaks at ∼x = 0 but is right skewed, while for genes in the GSR, the density peaks at ∼x = 7.5, showing that genes in the GSR are male-biased.

PGR has been proposed as a mechanism to reduce conflict between the somatic and germline genomes during early embryogenesis. Bryant et al. (2016) identified that the genes eliminated during PGR are expressed throughout lamprey embryogenesis and found ontological overrepresentation of these genes in germline development and oncogenesis^4^. However, PGR is closely tied to PGC specification in sea lamprey since germ cells are those that do not undergo PGR. Thus, we hypothesized that genes in the GSR might play roles in both early embryogenesis as well as gonadal differentiation and/or development. To address this, we assessed global differences in expression of genes in the GSR vs. somatic genome during gonadal development across differentiated ovaries and testes sampled in early, mid, and late developmental stages as well as in undifferentiated larvae and prospective males prior to testicular differentiation (See Supplementary Fig. 1 and Supplementary Table 1). This revealed the surprising result that almost all of the genes in the GSR exhibit male-biased expression during gonad development (Figs. 1c, 1d, Supplementary Fig. 7), while genes in the somatic genome were, overall, equally likely to be expressed in the female or male gonad (as expected): i.e., the density of the male:female gene expression ratio peaks at x = 0 for genes in the somatic genome (Fig. 1d). A possible explanation for this observation could be that females do not have the same GSR as males, since the reference genome for sea lamprey was generated using sperm DNA. To examine this possibility, we aligned individual BAM files from both male and female gonad samples to the indexed superTranscriptome and annotation file using the Integrative Genome Viewer (IGV)^22^. This revealed 410 transcripts from female gonad samples that mapped to either known or novel exons in the GSR (Supplementary Fig. 8a–8b). This suggests that females harbour the GSR but that it exhibits very low gene expression in female gonads, perhaps due to hypermethylation.

The mechanism of sex determination in lampreys remains unknown, and may involve both genetic and environmental factors^18,23–26^. The single elongated gonad remains histologically undifferentiated for up to several years, and the differentiation process is asynchronous in females and males (see^18^). Ovarian differentiation occurs in the larval stage, following synchronized and extensive meiosis and oocyte growth. A few small oocytes may also appear in future males, but testicular differentiation does not occur until the onset of metamorphosis ∼2–3 years later, when resumption of mitosis in the remaining undifferentiated germ cells produces spermatogonia^23^. It also appears that some larvae may be capable of undergoing sex reversal to males following ovarian differentiation^24^. Thus, a suite of genes could be turned on to initiate testicular differentiation. Our data suggest that female sea lamprey gonads harbour the same GSR as males but, with the exception of some rRNA and ribosomal protein-coding genes, females exhibited very low expression of the GSGs (Supplementary Fig. 8a–8b). Male-biased sex ratios under conditions of high larval density or slow growth have led to suggestions that primary sex differentiation in lampreys is influenced by environmental factors^25,26^. Environmental factors that influence the activation or silencing of genes in the GSR could, at least partially, control sex determination. In this case, low expression of the GSGs would result in a phenotypically female lamprey, whereas high expression in late larval or early metamorphosing lamprey would produce a male.

### Somatic paralogues of GSGs are expressed differently than germline paralogues

We observed that many of the GSGs had duplicated copies: of the 163 GSGs, 92 were found to have one or more paralogous copies in the GSR (Supplementary Table 4) while 89 have putative paralogs in the somatic genome, suggesting that some of the GSGs may have been recruited to the GSR to play specific roles in gametogenesis. The somatic paralogues of the GSGs were found distributed throughout the entire somatic genome, on every chromosome except chromosome 49 (Fig. 2a, Supplementary Table 7). To assess whether the somatic paralogues of the GSR genes exhibit similar sex-biased expression, we selected one paralogous gene per genome (somatic and germline) and generated a heatmap to compare somatic vs. GSR expression of the paralogous genes (Fig. 2b). In keeping with the somatic-wide pattern (Fig. 1d), this demonstrated that the somatic paralogues of the GSR genes do not exhibit the same sex-biased expression (Fig. 2b).

**Fig. 2:**
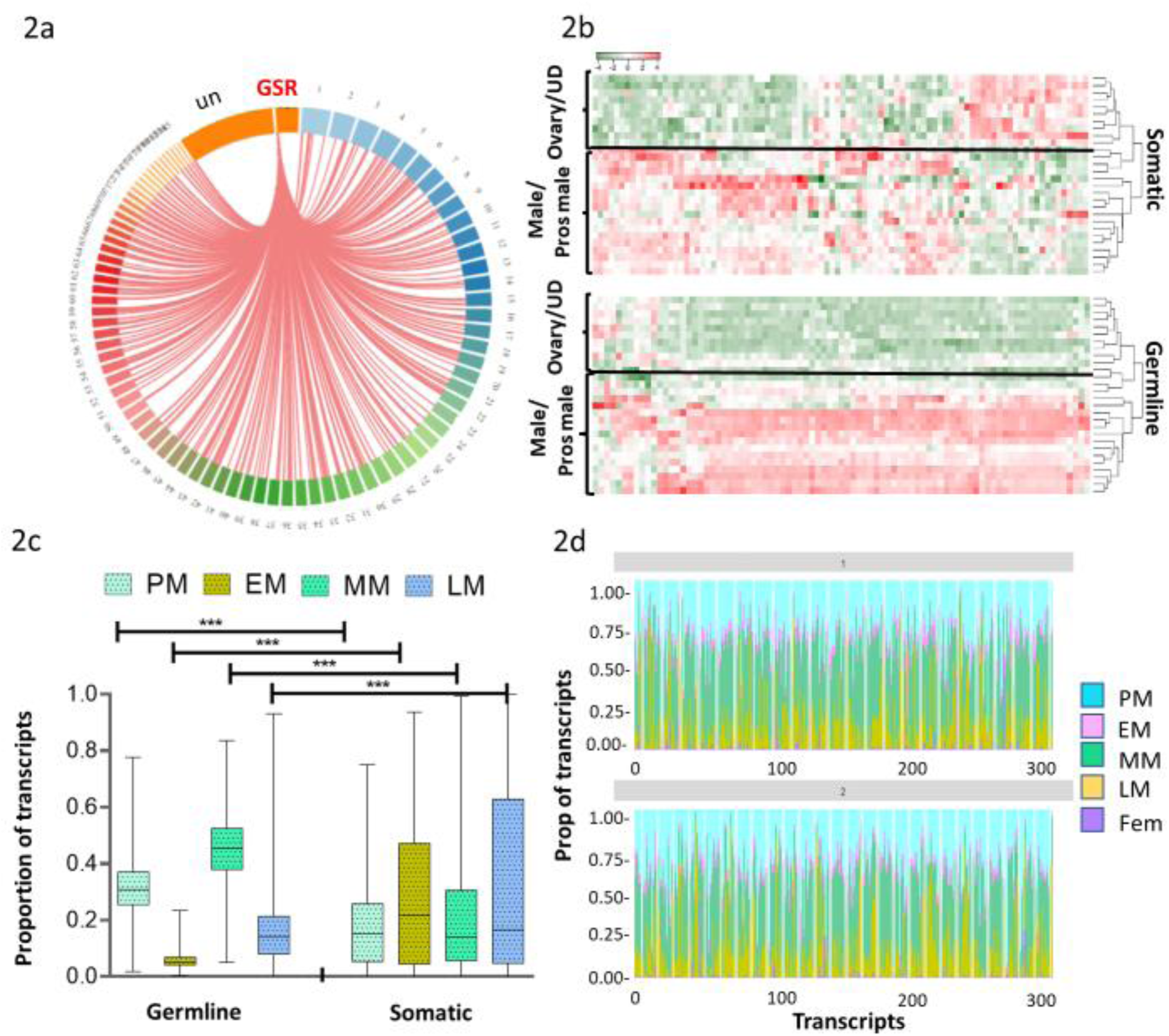
Somatic paralogues of GSGs are expressed differently than germline paralogues. **a)** Circos plot indicating the link between genes in the GSR with putative somatic paralogues in the sea lamprey genome. Chromosome 81 and enriched scaffolds are indicated as GSR and non-enriched scaffolds are indicated as un (unplaced scaffolds in somatic genome). **b)** Heatmap showing the relative expression of genes that have paralogues in both the GSR and somatic genomes in males and prospective males (pros male), females (ovary), and undifferentiated larvae (UD). **c)** Box plot showing the gene expression differences of somatic and GSR paralogues of GSGs in prospective males (PM), and early, mid, and late males (EM, MM, and LM, respectively). **d)** Comparison of the proportion of transcripts of GSGs in prospective, early, mid, and late males (PM, EM, MM, LM) and females (Fem). X-axis represents the number of transcripts present in GSR and Y-axis represents the proportion of transcripts in each stage.

Given the evidence of male-biased gene expression in the GSR, we next examined whether the GSGs had uniform expression across male gonadal developmental stages, using the number of male-biased genes per stage in the somatic genome as reference. In total, we identified 1270 male-biased genes (of 18,945 total genes), of which 409 (of 638 total genes) were found in the GSR and 861 (of 18,307 total genes) were found in the somatic genome, indicating that genes in the GSR have a 36x higher odds of exhibiting male-biased expression (OR = 36.5068 where *P* < 0.0001). Using the normalized counts of transcripts exhibiting male-biased expression, we compared the proportion of total transcripts in early, mid, and late testicular development and in prospective males by examining the interaction between genome (somatic or GSR) and stage using a repeated measures mixed model design in which gene nested in genome was a random effect, and stage was a repeated measure (Supplementary Table 8, Supplementary Fig. 9). This showed that there was a higher proportion of genes expressed in males in mid-testicular development and in prospective males in the GSR compared to somatic genomes, and a significantly lower proportion of genes expressed in early and late testicular development in the GSR relative to the somatic genome (Fig. 2c, Supplementary Tables 8 & 9).

To visualize the stage-specific bias in gene expression of GSGs, we plotted the relative proportion of transcripts expressed in each of the three male gonadal stages (early, mid and late), as well as in prospective males and the pooled sum of transcripts expressed at any female stage (Fig. 2d, Supplementary Table 10). This underscores that there is a similar pattern of expression across all genes in the GSR: high gene expression in prospective and mid gonadal stage males, but zero to very low expression in females. These findings are similar but distinct from those in zebra finch: the chromosomes undergoing chromosomal diminution and the genes eliminated are not sex-biased; however, individual genes have showed expression in both testes and ovaries, with overall greater enrichment for genes involved in ovarian development^16^. On the other hand, in a sciarid fly (*Sciara coprophila*), the elimination of one or two paternal X chromosomes in all somatic cells determines the sex of the embryo^10^. Here, we find evidence that the GSGs show comparatively higher expression in presumptive males when male sea lamprey are putatively undergoing sex determination and in a later stage of spermatogenesis when male gametes are generating spermatogonial Type B cells. This supports our hypothesis that gene expression in the sea lamprey GSR may function to control sex determination and/or differentiation, with high expression leading to testicular development in males and gene silencing resulting in ovarian differentiation in females.

Our IGV analysis showed that the GSR appears to be present in ovaries (Supplementary Fig. 8a-8b), suggesting that the genes in the GSR are turned off in females, while they are expressed in males throughout the sampled stages of spermatogenesis. One possibility is that differential DNA methylation is involved in sex determination/differentiation in sea lamprey. DNA methylation is a common process of epigenetic modification with known roles in gene regulation, embryogenesis and increasingly, sex determination^27^ which has, interestingly, become more important throughout deuterostome evolution^28^. A recent study in zebrafish (*Danio rario*) found that DNA methylation plays important functions in germline development as well as in sexual plasticity^29^. Given the clear role for the GSR in male spermatogenesis, we wanted to probe the expression of the GSR during early development bracketing PGR itself. To this end, we analyzed publicly available RNA-seq data from sea lamprey embryos that span the PGR (1–5 dpf). Of the 638 genes we identified in the GSR, only 186 were expressed during early embryogenesis. Of these 186 genes, 146 were expressed pior to PGR and 111 post-PGR, but only 20 had an average gene count >50 post-PGR and 18 pre-PGR (Supplementary Table 11), while the five most abundantly expressed genes code for ribosomal proteins. We then compared the expression of the 186 GSGs expressed during pre- and post-PGR embryos with our male and female gonad samples, and find that they exhibit very low expression in females and embryos, but high expression in male gonads (Supplementary Fig. 10). This further supports the hypothesis that the role of the GSR in sea lamprey is predominantly to support male gonadal development.

### Evolutionary conservation of GSGs and their function in vertebrate spermatogenesis

Genes in the GSR are expected to be released from the dosage sensitivity constraints of genes in the somatic genome^4^ and may show conservation of gene functions related to gonadal sex determination and differentiation in other vertebrates. We thus hypothesized that genes in the GSR 1) do not originate from a single linkage group in the pre-vertebrate ancestor, and do not map to a single linkage group in the post-2R vertebrate genome, 2) exhibit accelerated evolution either via high rates of duplication and/or amino acid change and 3) have known roles in sex determination or spermatogenesis in other vertebrates. To this end, we performed comparative mapping of genes in the sea lamprey GSR to an earlier chordate (*Branchiostoma belcheri*) and to nine post-2R taxa. Of the 163 unique gene names identified in the GSR, orthologues with variable levels of conservation across chordates were identified for 31 genes (Supplementary Table 12). Some of these genes are found predominantly as a single copy in most taxa, whereas in sea lamprey, we find a single copy in the GSR but multiple paralogues in the somatic genome (*rpab4, rlp37A, mid2bp*, and *hsop3*), multiple paralogues in both the GSR and somatic genomes (*scyp1*) or a single copy in the GSR and somatic genomes (*fgfr3*), or a single copy in the GSR but no copy in the somatic genome (e.g. *agrl3, cxb1, hsop3, rpab4*) (Supplementary Fig. 11, Supplementary Table 4).

On the other hand, some of the genes in the GSR are found in multiple paralogues in later vertebrate genomes, and in multiple copies in the GSR and/or somatic genomes in sea lamprey *(agrl3, cxb1, spop1, lpar1, cadh2, lrrn1, mlcl1*) (Supplementary Fig. 11). Of the 31 genes assigned to an orthogroup, 23 were also identified in *Branchiostoma* (Supplementary Table 12 and Supplementary Fig. 11), and 9 are present only in the GSR (not the somatic genome) suggesting that some of the lamprey GSR genes are not novel. Lastly, a comparative syntenic analysis between all genes in the lamprey genome for which orthogroups were assigned to the pre-vertebrate ancestral genome (n = 9,850) or human genome (n = 19,701) found blocks of conserved synteny for somatic but not GSGs. For example, there is strongly conserved synteny between the sea lamprey somatic genome and the 17 linkage groups hypothesized to exist in the pre-2R vertebrate genome (see ^29^), but the genes linked to the GSR are dispersed across all but two of the pre-2R linkage groups (Fig. 3a), and do not show conserved synteny in the human genome (Supplementary Fig. 12). This suggests that the genes involved in spermatogenesis in the GSR were independently duplicated into the GSR and were not part of an evolutionarily conserved paralogon.

**Fig. 3:**
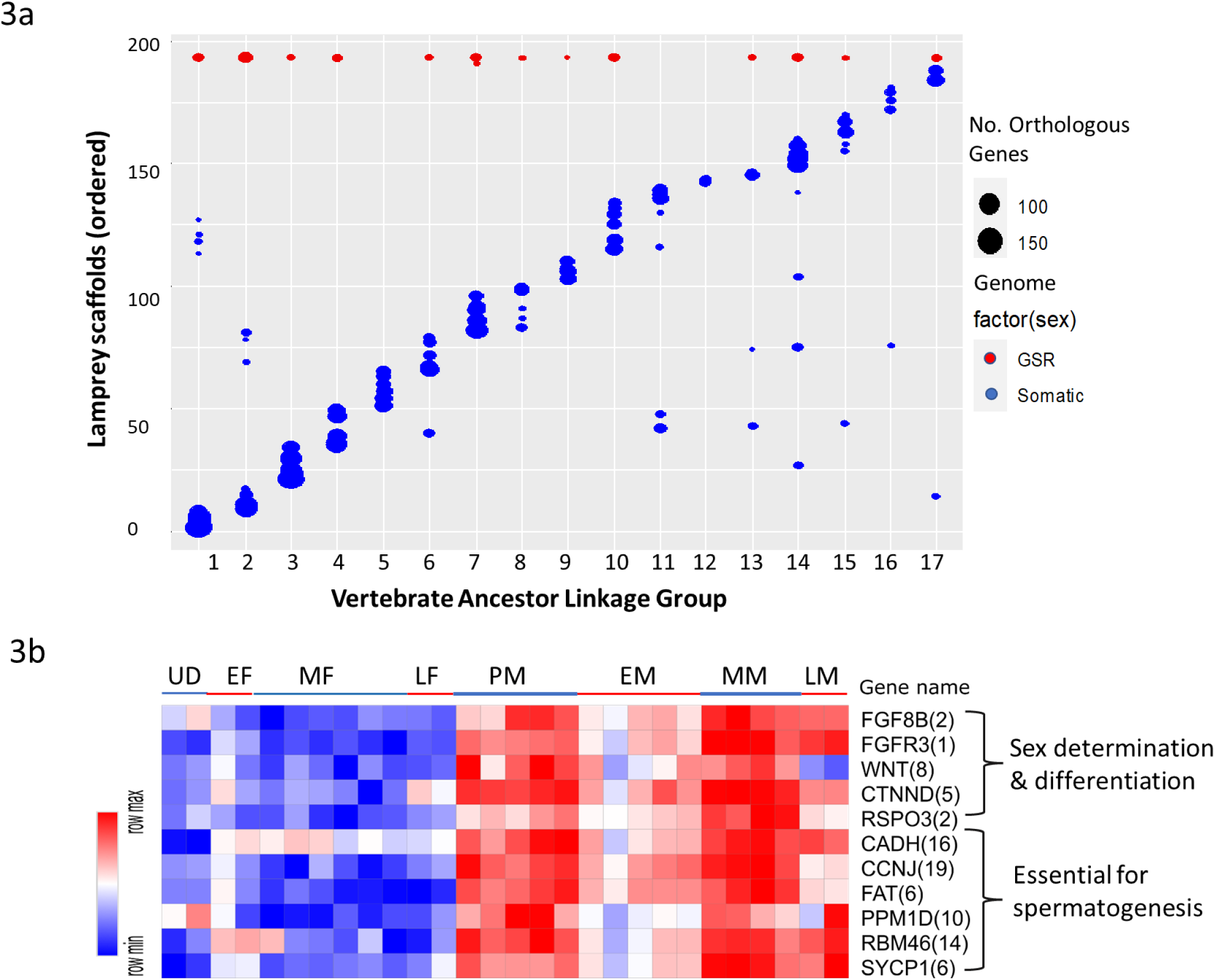
The evolutionary relationship of GSGs with the pre-vertebrate ancestral genome reconstructed by^30^ and and obtained from Genomicus webserver^31^ and their functional conserveness across vertebrate **a)** Chromosome plot showing comparative mapping in the sea lamprey and ancestral genomes. **b)** Heatmap of the median expression of paralogs in gene families present in GSR that have known roles in sex determination or spermatogenesis in other vertebrates. The numbers in brackets are the number of paralogues of these genes present in GSR. UD represents undifferentiated larvae, EF, MF and LF represent early, mid and late females respectively and PM, EM, MM and LM represents prospective, early, mid and late males respectively.

We searched the literature for evidence that any of the 163 unique gene names we identified in the GSR are associated with sex determination and/or differentiation in other taxa (Supplementary Table 13). Some of the GSGs have been found to exhibit female-biased expression in later diverging vertebrates, and some are involved in ovarian development, suggesting that the tissue (gonad) of expression may be conserved, but the function (male vs. female gonadogenesis) is not. Importantly, however, we find orthologues or paralogues of most of the core genes involved in sex-determination across vertebrates e.g., fibroblast growth factor 8 (*fgf8*), which is involved in sex determination in mice^32–34^, as well as fibroblast growth factor receptor 3 (*fgfr3*), which is involved in sex determination in sturgeon (*Acipenser dabryanus*)^35^. Other genes such as *scyp1*, which is important for early meiotic recombination during spermatogenesis^36^, R spondin (*rspo1)* and beta catenin 1 (*ctnnb1*) are important antagonists for Wnt pathway and initiating testicular differentiation^37,38^. Further, several of the gene families known to be essential for spermatogenesis are highly duplicated. For example, cadherins (*cadh*) are responsible for maintaining the integrity of testis structure^39^; cyclins (*ccnb*) are essential for cell progression during distinct phases of the male spermatogenesis pathway ^40^; RNA binding proteins (*rbm*) play diverse and important roles in spermatogenesis including testis-specific splicing^41^ and the absence of *rbm46* (present in 16 copies in the sea lamprey GSR) is associated with male infertility in mice^42^. Other important genes e.g., *sox9* and *cbx2* which play roles in stabilizing the male differentiation pathway, are present in the somatic genome of sea lamprey. We find that all of these genes are highly expressed in the gonads of prospective males and mid-males when gonadal germ cell specification and spermatogonial development are occurring respectively (Fig 3b). This suggests that the GSR is likely to play a role in gonadal sex determination and differentiation as well as spermatogenesis in sea lamprey.

### Phylogenetic relationship of GSGs provides evidence of diversified genes involved in sex-determination pathway

Lampreys diverged from the jawed vertebrate lineages more than 500 million years ago^43,44^, either after the two rounds (2R) of whole genome duplication (WGD) that occurred in early vertebrate evolution^30,45^, or more likely after 1R^20,46,47^. However, a recent study suggested that, after the 1R tetraploidization, lampreys underwent an additional hexaploidization^48^. Since lampreys have an unusual vertebrate ploidy state, it proved impossible to perform a reliable test of positive selection at the amino acid level (which requires essentially gapless alignments) for the germline genes in agnathans (lampreys and hagfishes) relative to other vertebrates. Thus, we selected a few genes which have important roles in gametogenesis in other species for phylogenetic analyses (see Supplementary Table 13).

Gene trees were reconstructed using the output from OrthoFinder and the orthologues of sea lamprey GSGs in 10 other chordates identified (see Supplementary Fig. 11) and combined with our data on gene annotations and genomic location (GSR vs. somatic) in sea lamprey. This revealed that the *cadh* gene family is highly duplicated in both the germline and somatic genomes of sea lamprey (16 vs. 15 duplicates, respectively) andhagfish (Supplementary Table 12). In particular, *cadh2* has undergone a divergent expansion in the GSR in sea lamprey (Supplementary Fig. 13a); agnathans have witnessed an expansion of a somatic cluster of genes related to vertebrate *cadh1/cadh3/cadh13* as well as an expansion in both the somatic and germline genomes of a novel cadh paralogue (bottom of Supplementary Fig. 13a). Phylogenetic trees for *hykk* (Supplementary Fig. 13b), *sycp1* (Supplementary Fig. 13c), and *adgrl* (Supplementary Fig. 13d) depict similar patterns of one or more highly duplicated germline lineages that are sometimes interspersed with closely related somatic paralogues (*hykk* and *scyp1*), but overall, they show clades of highly diversified germline lineages marked by long internal branch lengths, indicating that the GSGs exhibit independent evolution for variable periods of time and may be subject to positive selection.

We identified a novel *fgfr3*-like gene in the germline genome, and confirmed expression of a possible ligand for it, *fgf8b*, also located in the germline genome. The *fgfr3*-like gene was not identified by OrthoFinder as an orthologue of the somatic copy of *fgfr3*; thus, we downloaded the canonical coding sequences for *fgfr3*, and a related gene also present in the somatic genome, *fgfrl1*, from eight post-2R taxa and reconstructed a ML tree with bootstrap support (Fig. 4a). This revealed that the germline sequence of the *fgfr*-like coding sequence is more closely related to *fgfr3* in the sea lamprey somatic genome and to the *fgfr3* in higher vertebrates (bootstrap support 100%), while the somatic copy of *fgfrl1* groups with the *fgfrl1* sequences from later vertebrates and there is no paraloge in the GSR (100% bootstrap support). Examination of the expression of these three genes as well as the possible receptor for the germline gene, *fgf8b*, indicates that the germline copy of *fgfr3* and *fgf8b* have very low expression in female gonads, and somewhat higher expression in male gonads: notably, *fgf8b* is most highly expressed in prospective male and mid-stage male gonads (Fig. 3b, 4b–4e). Given their role in sex determination in other vertebrates, sea lamprey germline genes *fgf8b* and *fgfr3* warrant further investigation as possible loci involved in sex determination.

**Fig. 4:**
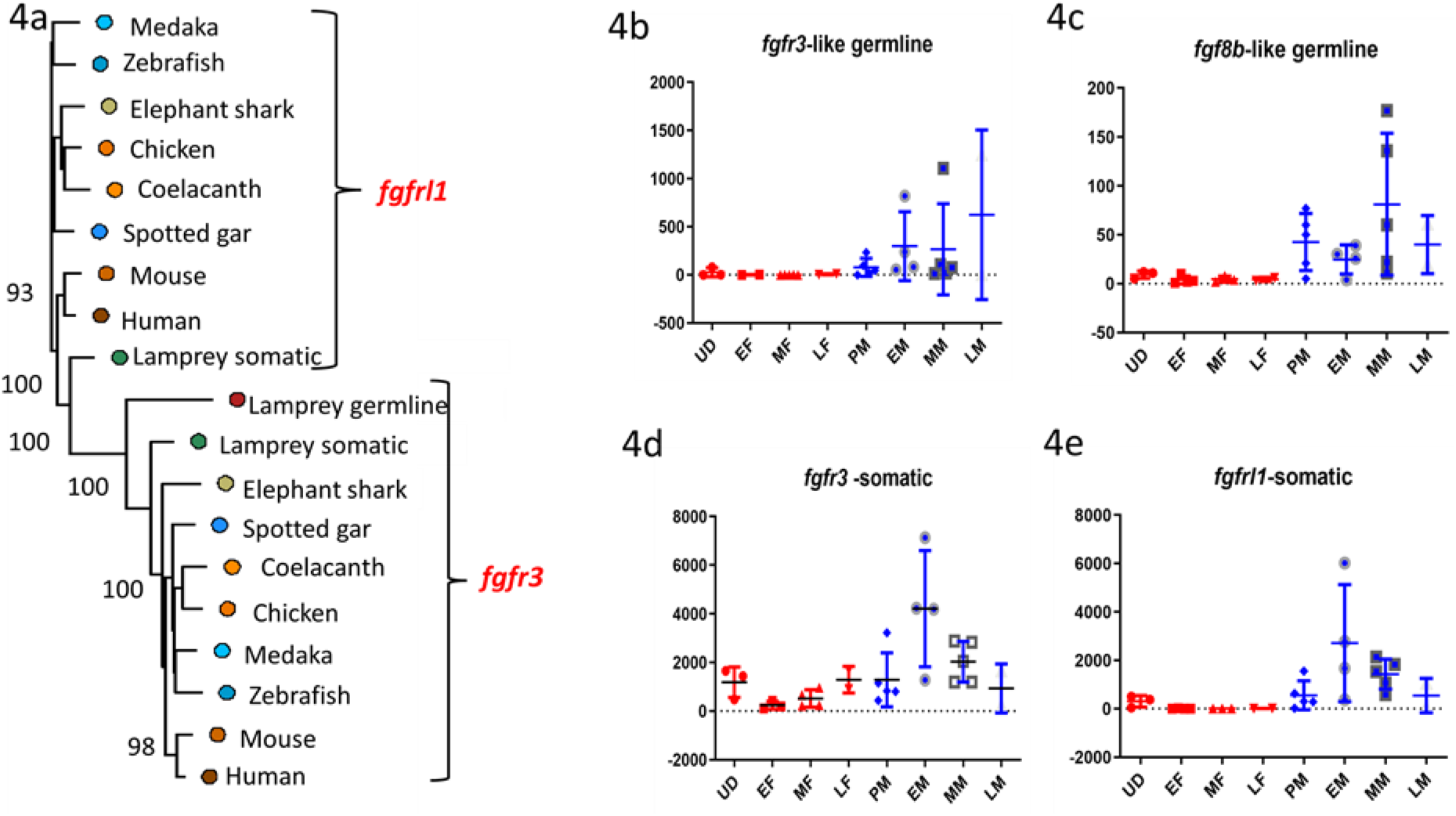
Phylogenetic tree of putative ligand and receptor of fibroblast growth factor (fgf) and their genome-, sex- and stage-specific expression a) phylogenetic tree, b) *fgfr3*-like gene in the GSR genome, c) *fgf8b* like gene in GSR, d) *fgfr3* in somatic genome, and e) *fgfrl1* in the somatic genome. In 4b–4e, the Y-axis is gene counts and the X-axis is stage, where UD is undifferentiated larvae; EF, MF, and LF are early, mid, and late females; and PM, EM, MM, and LM are prospective, early, mid, and late males, respectively.

## Conclusion

The study of PGR events and their effects on gonadal development and sex determination represent a burgeoning field in evolutionary biology. Our result suggests that the genes present in the GSR in sea lamprey are involved in the crucial processes of sex differentiation and testicular development, and could be involved in sex determination. We find GSGs are most highly expressed in prospective males and in males undergoing spermatogonial differentiation, but they have low overall expression in females. Given evidence that sex is partially determined by environmental factors in sea lamprey, the possible role of methylation in the GSR during early stages of gonad development in larval sea lamprey warrants further attention.We find low levels of syntenic or sequence conservation of genes in the GSR across chordates, but importantly, many of the genes identified in the GSR are known to play roles in gonad differentiation or sex determination in other vertebrates. Assuming females harbour the same GSR as males, our data suggests that the factors controlling epigenetic modification of the GSR are pivotal for sex determination and differentiation. Further work is needed to assess the presence and chromatin accessibility of the GSR in females and to identify the function of the GSGs in sea lamprey sex determination and differentiation.

## Methods

### Sample preparation and RNA extraction

Sea lamprey from different life-history stages were collected by collaborators using these samples for other projects. An Abbreviated Protocol for Minimal Animal Involvement form completed at the University of Manitoba determined that an Animal Use Protocol (AUP) was not required because live sea lamprey were not handled by us for the purposes of this project, and no animals were sacrificed or manipulated solely to provide us with tissue.

Larval sea lamprey were collected by backpack, pulsed DC electrofishing in tributaries of the Richibucto River, New Brunswick, Canada, or in tributaries of Lake Huron and Lake Michigan in the Great Lakes basin (Supplementary Table 1). Larvae were transported or shipped live to Wilfrid Laurier University, Waterloo, ON, sorted according to size, and transferred to 110 L holding tanks supplied with aerated well water at a flow rate of 1.0–2.0 L/min. The larvae were monitored for external signs of metamorphosis (e.g., changes in eye and oral disc morphology) and then euthanized at the desired stages. The brain and gills, required for other projects, were dissected and placed in RNAlater. With the remaining carcass (posterior to the last branchial pore), RNAlater was injected into the gut to perfuse the intestine, liver, gallbladder, kidneys, and gonad. The carcass with organs was then placed in a 10 mL Falcon tube and filled with RNAlater to saturate the tissues thoroughly. Dissections were completed as rapidly as possible to reduce any potential RNA degradation. Samples were kept at 4 °C for 24 h, stored at –80 °C, and then shipped to the University of Manitoba on dry ice, and stored at –80 °C upon arrival. The gonads were subsequently dissected out and placed in a 1.5 mL centrifuge tube with 1 mL RNAlater and kept at –20 °C. Sex was identified during dissection based on physical inspection with the naked eye (i.e., the ovary is larger and has a different texture than the testis or undifferentiated gonad), and gonadal stage was identified by a combination of visual inspection and inferences based on larval size and stage of metamorphosis^18^ (Supplementary Table 1).

Adult sea lamprey were captured in traps near the mouth of the Black Mallard River or Ocqueoc River, MI, during their upstream (spawning) migration (Supplementary Table 1).Lamprey were euthanized, length and weight measurements were taken, and ∼35 mg gonad was flash frozen in a 2.0 mL centrifuge tube and kept on dry ice (April 2018) or placed in a 1.5 mL centrifuge tube with 1 mL RNAlater and kept at –20 °C (June 2018). Samples were shipped to the University of Manitoba on dry ice, and stored at –80 °C.

Total RNA was isolated from ∼30 mg of gonadal tissue from each individual using the RNeasy Mini kit (Qiagen, USA) according to the manufacturer’s protocol. The extracted RNA was treated with RNase-free DNase set (Qiagen, USA) to remove residual genomic DNA. RNA quantity and quality was assessed using a NanoVue Plus spectrophotometer. The RNA samples were preserved at –80 °C.

To obtain a comprehensive representation of gene expression, RNA from individuals at the same stage of development and same sex was pooled. Early males (n = 4) were those identified by external morphological characteristics to be in the early to mid stages of metamorphosis and thus presumed to be in the early stages of spermatogonial differentiation, that is, in the process of producing Type A spermatogonia (Supplementary Fig. 1, Supplementary Table 1)^18^. Mid males (metamorphosing stage 7 and immediately post-metamorphosis; n = 6) were presumed to be undergoing spermatogonial proliferation and differentiation and producing Type A and Type B spermatogonia, while late males were sexually mature (n = 2). In early females (n = 2), ovarian differentiation had been initiated and/or completed (i.e., with a number of small growing oocytes in the gonad), mid-stage females (n = 6) had completed oocyte differentiation and were arrested in meiotic prophase with larger growing oocytes, and late females were sexually mature (n = 2). In addition to samples that were definitively male and female, larvae that had histologically undifferentiated gonads and were below the size at which ovarian differentiation occurs (n = 2) and presumptive male larvae with histologically undifferentiated gonads but beyond the size at which ovarian differentiation is complete (n = 4) were included.

### Library preparation, Illumina sequencing, and data filtering

High-quality RNA from 28 gonad samples was sent to Genome Quebec, McGill University, Montreal, to construct a cDNA library and perform RNA sequencing. Messenger RNA (mRNA) was isolated using poly-A isolation and non-normalized libraries prepared using the Illumina TruSeq DNA Kit and Epicentre Script Seq Kit. Sequencing was performed in both forward and reversed directions and 100 base pair (bp) reads were generated on an Illumina Hi-Seq 4000 PE100. The resulting RNA-Seq paired-end (PE) reads were checked for quality control using FASTQC (v0.11.8)^49^, and low-quality sequences and adapters were trimmed with Trimmomatic (v0.36)^50^, using ILLUMINACLIP:TruSeq3-PE-2.fa:2:15:10 LEADING:5 TRAILING:5 SLIDINGWINDOW:4:5 MINLEN:50 and a quality score threshold of Phred-33.

### Combining reference and *de novo* assemblies

#### Generating comprehensive gonadal superTranscriptome

The software pipeline Necklace^19^, was used to generate a merged superTranscriptome derived from three sources: 1) a genome-guided alignment using the sea lamprey reference genome, 2) a *de novo* assembly using Trinity, and 3) a reference-based proteome from other chordate species. For the genome-guided assembly, the 28 gonadal transcriptomes were mapped to the Vertebrate Genome Project (VGP) sea lamprey reference germline genome (https://ftp.ncbi.nlm.nih.gov/genomes/all/GCF/010/993/605/GCF_010993605.1_kPetMar1.pri/GCF_010993605.1_kPetMar1.pri_genomic.fna.gz) and associated gene annotation file (https://ftp.ncbi.nlm.nih.gov/genomes/all/GCF/010/993/605/GCF_010993605.1_kPetMar1.pri/GCF_010993605.1_kPetMar1.pri_genomic.gff.gz) available at NCBI. Reads were aligned to the sea lamprey genome using HISAT2, and StringTie^51^ was used to assemble transcripts, some of which map to known genes and some of which are novel (MSTRG IDs). For the third tier of the Necklace pipeline, reference proteomes from a non-teleost fish, spotted gar (*Lepisosteus oculatus*) (https://ftp.ncbi.nlm.nih.gov/genomes/all/GCF/000/242/695/GCF_000242695.1_LepOcu1/GCF000242695.1_LepOcu1_protein.faa.gz), and a cartilaginous fish, elephant shark (*Callorhinchus milii*) (https://www.ncbi.nlm.nih.gov/genome/689?genome_assembly_id=49056) were used.

In the second step, a *de novo* assembly of reads was generated with Trinity^52^ for all 28 samples. The assembled transcripts from genome-guided and *de novo* assembly were sorted into three groups: annotated transcripts that align to the reference genome (known genes), transcripts that align to the reference genome but are not found in the reference annotation (reference-based novel genes), and unmapped novel transcripts – those that align to the spotted gar/elephant shark proteome (*de novo-*specific genes). These three groups were merged into a single superTranscriptome and used for the second stage of the analysis: gene counting and differential expression analyses. The Necklace pipeline allows for the identification of novel transcripts yet generates a compact and comprehensive superTranscriptome, while preventing the introduction of false chimeras generated during *de novo* assembly. The step-by-step workflow of Necklace pipeline is illustrated in Supplementary Fig. 14.

In total, we identified 42,479 genes in the sea lamprey germline genome, of which 20,630 overlapped with those annotated by NCBI (representing ∼94% of the total number of genes in the VGP annotation), 21,808 were identified *de novo* through StringTie, and 40 Trinity *de novo* assembled transcripts matched sequences in the spotted gar/elephant reference proteome by homology. However, since the genomic location of these 40 homology-based sequences could not be ascertained, they were discarded from further analyses. Of the remaining 42,439 sequences, tRNA, rRNA, and lncRNAs (long non-coding RNAs), were removed, retaining 18,945 protein-coding transcripts (16,328 from the VGP annotation and 2,617 novel transcripts, which is ∼14% of the total gene list) (Supplementary Fig. 15). Those 18,945 genes pertain to 12,583 unique gene names, which would be a lower limit on the actual number of genes identified, since paralogous genes may be assigned the same gene name.

#### Gene-counts

Reads from each of the 28 gonadal transcriptomes were subsequently aligned to the merged superTranscriptome, and gene counts extracted and filtered. These gene-counts are used for further downstream analysis, i.e., in differential gene expression analysis, identifying sex-biased and sex-specific transcripts and genes.

### Functional annotation and identifying orthogroups

#### Functional annotatio

All of the 18,945 putatively protein-coding genes generated from the Necklace pipeline were annotated using Trinotate pipeline (v3.2.0)^53^ following the method described at (http://trinotate.github.io/). Initially, Transdecoder (v5.5.0) was used to obtain the expected start and stop sites of protein translation from the assembled superTranscriptome. Then each transcript and protein sequence were searched against the SwissProt database using blastx and blastp. The HMMER algorithm was used to search PFAM (ran in hmmer (v3.2.1)) for protein domain identification, signalp (v4.1f) and tmHMM (v2.0c) were used to predict the signal peptide and transmembrane regions, respectively, and Rnammer (v1.2) was used to identify rRNA transcripts which were automatically removed in a later stage of the pipeline. In the final stage, the results from blast searches were combined with the other functional annotation data and loaded into the Trinotate.SQLite database: an e-value of 1e-5 was used as the threshold to generate the functional annotation report.

#### Orthogroup identification

The homology between the genes in our annotated sea lamprey gonadal superTranscriptome were compared to genes in 11 chordate species chosen to represent important time points in chordate evolution using the OrthoFinder pipeline^54^. Protein sequences were obtained from human (ftp.ensembl.org/pub/release-103/fasta/homo_sapiens/pep/Homo_sapiens.GRCh38.pep.all.fa.gz), mouse (*Mus musculus*) (ftp://ftp.ensembl.org/pub/release-102/fasta/mus_musculus/pep/Mus_musculus.GRCm38.pep.all.fa.gz), zebrafish (ftp.ensembl.org/pub/release-103/fasta/danio_rerio/pep/Danio_rerio.GRCz11.pep.all.fa.gz), chicken (ftp.ensembl.org/pub/release-103/fasta/gallus_gallus/pep/Gallus_gallus.GRCg6a.pep.all.fa.gz), medaka (*Oryzias sinensis*) (ftp.ensembl.org/pub/release-103/fasta/oryzias_sinensis/pep/Oryzias_sinensis.ASM858656v1.pep.all.fa.gz), spotted gar (ftp.ensembl.org/pub/release-103/fasta/lepisosteus_oculatus/pep/Lepisosteus_oculatus.LepOcu1.pep.all.fa.gz), elephant shark (ftp.ensembl.org/pub/release-103/fasta/callorhinchus_milii/pep/Callorhinchus_milii.Callorhinchus_milii-6.1.3.pep.all.fa.gz), coelacanths (ftp.ensembl.org/pub/release-103/fasta/latimeria_chalumnae/pep/Latimeria_chalumnae.LatCha1.pep.all.fa.gz/), hagfish (*Eptatretus burgeri*) (ftp.ensembl.org/pub/release-103/fasta/eptatretus_burgeri/pep/Eptatretus_burgeri.Eburgeri_3.2.pep.all.fa.gz), amphioxus (*Branchiostoma belcheri*) (https://ftp.ncbi.nlm.nih.gov/genomes/all/GCF/001/625/305/GCF_001625305.1_Haploidv18h27/GCF_001625305.1_Haploidv18h27_protein.faa.gz). OrthoFinder uses the complete list of known protein sequences from all included taxa to find putative orthologues, and then creates orthogroups with related sets of orthologues. OrthoFinder exports multiple sequence alignments and rooted gene trees for all orthogroups, which can be used to infer gene duplication events. Overall, in this study, 93.1% of the genes in the 12 chordate species were assigned to one of 27,364 orthogroups, and 5606 orthogroups contained representatives of all 12 species.

### Prediction of germline-specific region (GSR) and genes by enrichment analysis

#### Identifying GSGs in the GSR

Although the GSR in sea lamprey has been identified for a previous germline assembly (gPmar100)^20^, when the new VGP germline genome was deposited on NCBI, the corresponding positions were not available. Following the protocol from Smith et al. 2018, germline enrichment was calculated using the DifCover program by calculating differences in read depth between a single germline sample (sperm) and a single somatic sample (blood) from the same male. We downloaded and mapped the same sperm (SRR5535435) and blood (SRR5535434) samples they had used from their previous analysis to identify the GSR coordinates in the newly-deposited VGP genome in order to facilitate our downstream transcriptomic analyses. The DNAcopy output file was generated by following step by step workflow with default settings described in the DifCover pipeline (https://github.com/timnat/DifCover) (See Supplementary Fig. 14)^21^. This DNAcopy output file was then used to identify GSR from the new chromosome level assembly, VGP germline genome and the DNAcopy output file. Later, the GSGs were identified by extracting all genes that fell within regions having an enrichment score >2 using bedtools (v2.29.0) with the aid of the genome-based annotation file (generated in Necklace pipeline).

Initially, we identified 1845 GSGs by extracting the DNAcopy output file from VGP genome; however, only 783 protein-coding GSGs were retained with gene counts after the initial filtration steps discussed in previous section. In the next step, we sorted genes based on their location: if two genes with the same name had overlapping start and end points, the canonical transcript was retained, which reduced the number of genes to 672. In the final step, we extracted the protein sequences associated with each of these genes from the transdecoder pep file and removed ambiguous sequences. The final list consisting of 638 GSGs was merged with the Trinotate annotation report to assign putative gene names for the novel genes, and with the reference annotation for genes identified by VGP. In total, 163 unique gene names were assigned to the 638 GSGs, 70 of them in a single copy, and the remaining 93 in 2–77 copies (Supplementary Fig. 2).

Given our finding that genes in the GSR are highly expressed during gonad development (see Results and discussion), we wanted to assess whether all or a subset of the genes in the GSR are also expressed during early embryonic development. To this end, we downloaded paired-end RNA-seq read data for embryos sampled at 1 dpf (SRR3002837), 2 dpf (SRR3002840), 2.5 dpf (SRR3002843), 3 dpf (SRR3002846), 4 dpf (SRR3002849) and 5 dpf (SRR3002852) from the SRA database (https://www.ncbi.nlm.nih.gov/sra?linkname=bioproject_sra_all&from_uid=306044). Reads were aligned to the VGP genome using HISAT2 (v 2.2.1)^51^, and assembled into transcripts, and gene and transcripts counts were obtained per sample using Stringtie (v 2.0)^51^. In the next step, we extracted the embryo-expressed GSGs using the merged annotation file generated in the previous step and the DNACopy output file generated from DifCover analysis (Supplementary Fig. 13). We considered only genes available in the reference annotation that mapped to our GSR, which resulted in 184 GSGs (after filtering ncRNA (non-coding RNAs), rRNA (ribosomal RNAs) and pseudogenes from the reference annotation) expressed in early embryonic development of which 149 genes overlapped with those expressed in our gonad samples. The gene count file was used to compare gene expression in the GSR pre-PGR (1dpf, 2 dpf and 2.5 dpf) and post-PGR (3 dpf, 4 dpf and 5 dpf) and between gonads and embryos (both pre-PGR and post-PGR).

#### Identifying somatic copies of GSGs

To identify putative paralogues of GSGs in the somatic genome, the list of all genes was sorted based on the unique gene names obtained from either the Trinotate annotation report or from the reference annotation. Of the 163 unique genes identified in the GSR, 89 were found to have either a single or multiple putative paralogues in the somatic genome, of which 31 were matched to a unique OrthoGroup by OrthoFinder (Supplementary Fig. 16).

### Identifying male-biased expression

To identify sex-biased genes, only the gonadal transcriptomes from definitive males were analyzed; the undifferentiated larvae and prospective males were removed from this comparison. Since traditional differential gene expression analyses using a threshold log-fold change between conditions would likely identify genes with low or no expression in one sex but also low expression in the other sex, we aimed to target genes that were very highly expressed in one sex but had much lower or no expression in the other. The gene count data was obtained from the raw gene count data across samples with reference to the necklace superTranscriptome, and these were later filtered using normalized gene counts and log fold change (logFC) using DESeq2^55^ and EdgeR^56^. That gave us an estimate of 6088 genes with higher logFC in males than females of which top 20% genes were considered to be male-biased as long as the total gene count is equal or more than 1000 (Supplementary Fig. 17).

### Comparison of GSG expression across sex and stage and functional enrichment analysis

#### Comparison of GSG expression across sex and stage

The genome-wide raw gene counts were converted to normalized counts using DESEQ2^55^, and the log2 expression of all genes in the GSR compared in a sex- and stage-specific manner. To assess global differences in sex bias in gene expression, we compared the density of the relative log2(male:female normalised expression) of all genes in the somatic genome vs GSR. To compare the difference in expression between GSGs against their somatic paralogues across both sexes and stage, we calculated the mean normalized log2 gene count and visualised the data with a heat map. To assess whether GSGs exhibit stage-specific sex-biased expression, we extracted the list of all genes exhibiting sex-biased expression (as defined above) in both the somatic genome and GSR, and assessed whether the proportional expression of genes differed between genomes and stage using a repeated measures mixed model in which proportion gene expression was the response and the model was gene(genome) + stage + genome*stage, with gene as a random effect, and stage as the repeated measure.

#### Functional enrichment

The list of genes identified in the GSRs was submitted for pathway analysis using the human protein-coding genes as background using PANTHER (v14) (http://pantherdb.org) ^57^. The PANTHER GO-slim molecular process terms associated with each gene were used for an over-representation test^57^ in which the Fisher exact test was performed to assess the significance of terms at an FDR of 0.05. Additionally, we used the gene ontology (GO) terms associated with the GSGs identified by Trinotate^53^, and visualized them in REVIGO (http://revigo.irb.hr/)^58^ in a scatter plot that shows cluster representatives in a two-dimensional space derived by GO terms with semantic similarity measure and clustering set at 0.9 overall. Terms were plotted with size proportional to fold-enrichment above expected and color according to the log10 of the FDR p-value (Fig. 1b; see Results and discussion).

### Comparative mapping and phylogenetic analysis

#### Comparative mapping

Lampreys, being an intermediate lineage between 1R and 2R WGD, are important model organisms for the study of evolution of genes as well as evolution of physiological process^20,30,45–48^. We compared the evolutionary origin and relationship of genes in the GSR to the pre-2R vertebrate genome and in later diverging taxa. To this end, we identified orthologues of the genes from all 85 assembled chromosomes as well as to scaffolds that we identified as enriched (GSR) or non-enriched (somatic) for germline DNA (see results) to those in human, chicken, and spotted gar and a reconstructed pre-2R vertebrate genome. Orthologous genes were identified using the output of OrthoFinder^54^, assigned to their chromosomal location using BioMart (ENSEMBL) and the number of co-orthologues per linkage group/chromosome calculated pairwise. To remove marginally supported syntenies, we used the observed number of lamprey orthologues identified by chromosome for each species comparison as the maximum expected value and retained all syntenic chromosomal pairs; if there were >10 genes shared between species for lamprey chromosomes 1–69, or > 5 for chromosomes 70–85 (this criterion was set based on chromosome size), and retained all orthologous gene matches for the GSR. The reconstructed pre-2R vertebrate genome was downloaded from the ftp site of the Genomicus webserver^31^ (ftp://ftp.biologie.ens.fr/pub/dyogen/genomicus/69.10 details of the reconstruction are described in^30^).

#### Phylogenetic analysis

Given that genes in the GSRs may have unique evolutionary histories, the phylogenetic relationships of a subset of the genes in the GSRs and their somatic orthologues were reconstructed along with orthologous/paralogous genes identified from the 11 taxa included in the OrthoFinder output. Phylogenetic trees were obtained from OrthoFinder which uses RaxML reconstruction^54^. Trees were not available for all GSGs, including the sea lamprey putative orthologue of *fgfr3*, which has been shown to be important for sex determination and differentiation in other taxa^32,35,59,60^. Thus, we obtained sequences for *fgfr3* for the same 11 species employed in the OrthoFinder analyses, and then performed an alignment in Maaft (https://mafft.cbrc.jp/alignment/server/) followed by ML reconstruction with RAXML. We hypothesized that genes in the GSR are under relaxed evolutionary constraint and relaxed dosage sensitivity and thus may exhibit accelerated rates of sequence evolution. However, we were unable to employ tests of dN/dS due to the difficulty of obtaining sufficiently un-gapped alignments of the coding sequence of the sea lamprey genes relative to those from other jawed vertebrates. Nevertheless, phylogenetic trees were generated to understand the relationship of paralogous copies of the GSGs in the GSR to those in the somatic genome, as well as the relationship of the protein coding sequences in sea lamprey to those in other chordate taxa.

## Data availability

The RNA-sequencing reads used for this study have been deposited in the NCBI repository under the BioProject accession number PRJNA749754 and will be available upon publication of manuscript.

## Acknowledgement

We thank Dr. John B. Hume (Michigan State University), Dr. Nicholas Johnson (U.S. Geological Survey), Dr. Michael Wilkie (Wilfrid Laurier University), and Joshua Sutherby (University of Manitoba) for providing us with the samples used in this study. We also thank Arfa Khan (University of Manitoba) for her guidance during sample processing and RNA extraction.

