## Supplementary Figures for "The germline-specific region of the sea lamprey genome plays a key role in spermatogenesis"

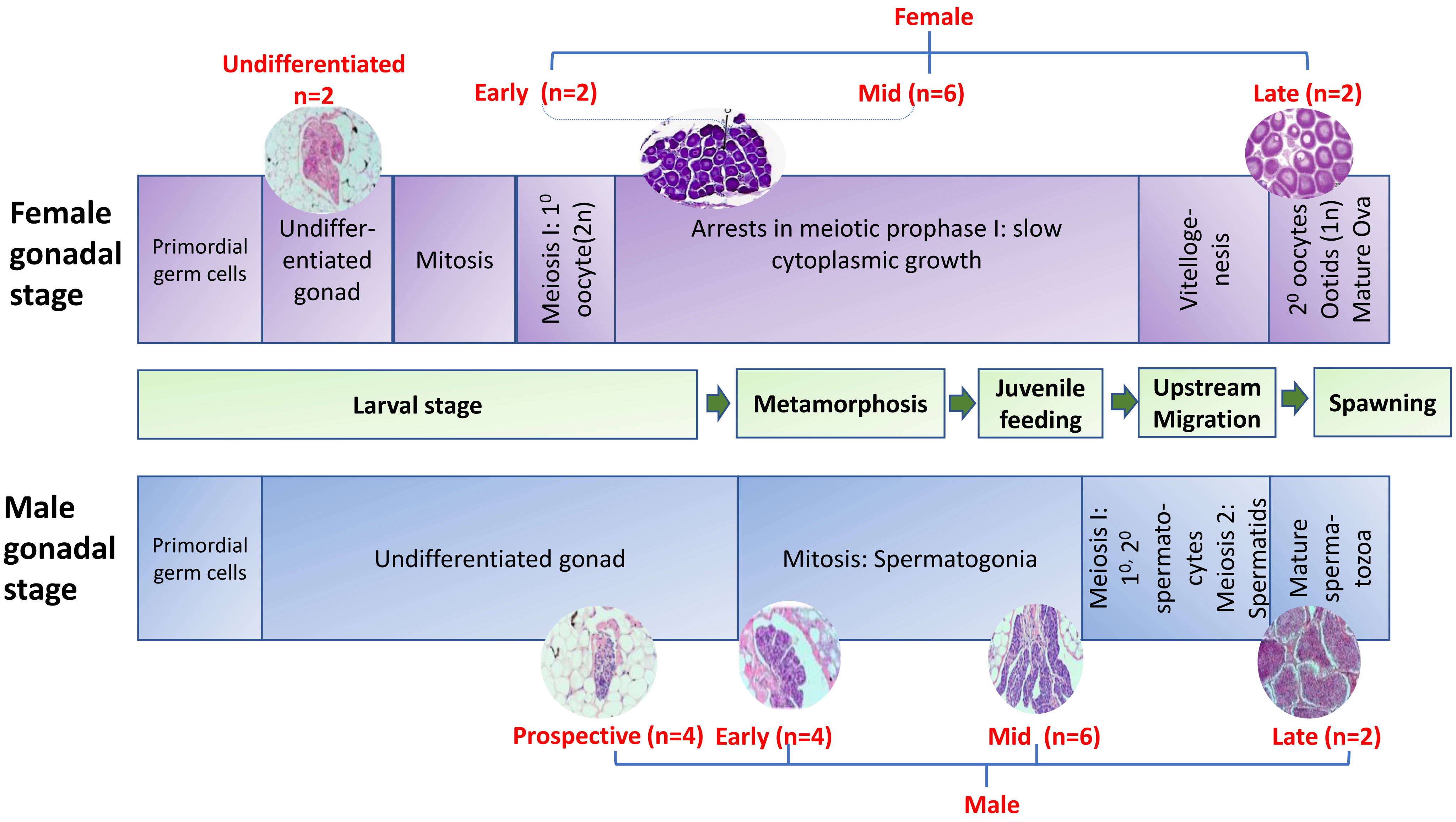


**Supplementary Fig. 1:** Schematic presentation of sea lamprey gonadal and life-history stages examined in this study; sample collection details are provided in Supplementary Table 1. The figure is adapted from Docker et al. (2019), with images from Khan (2017).

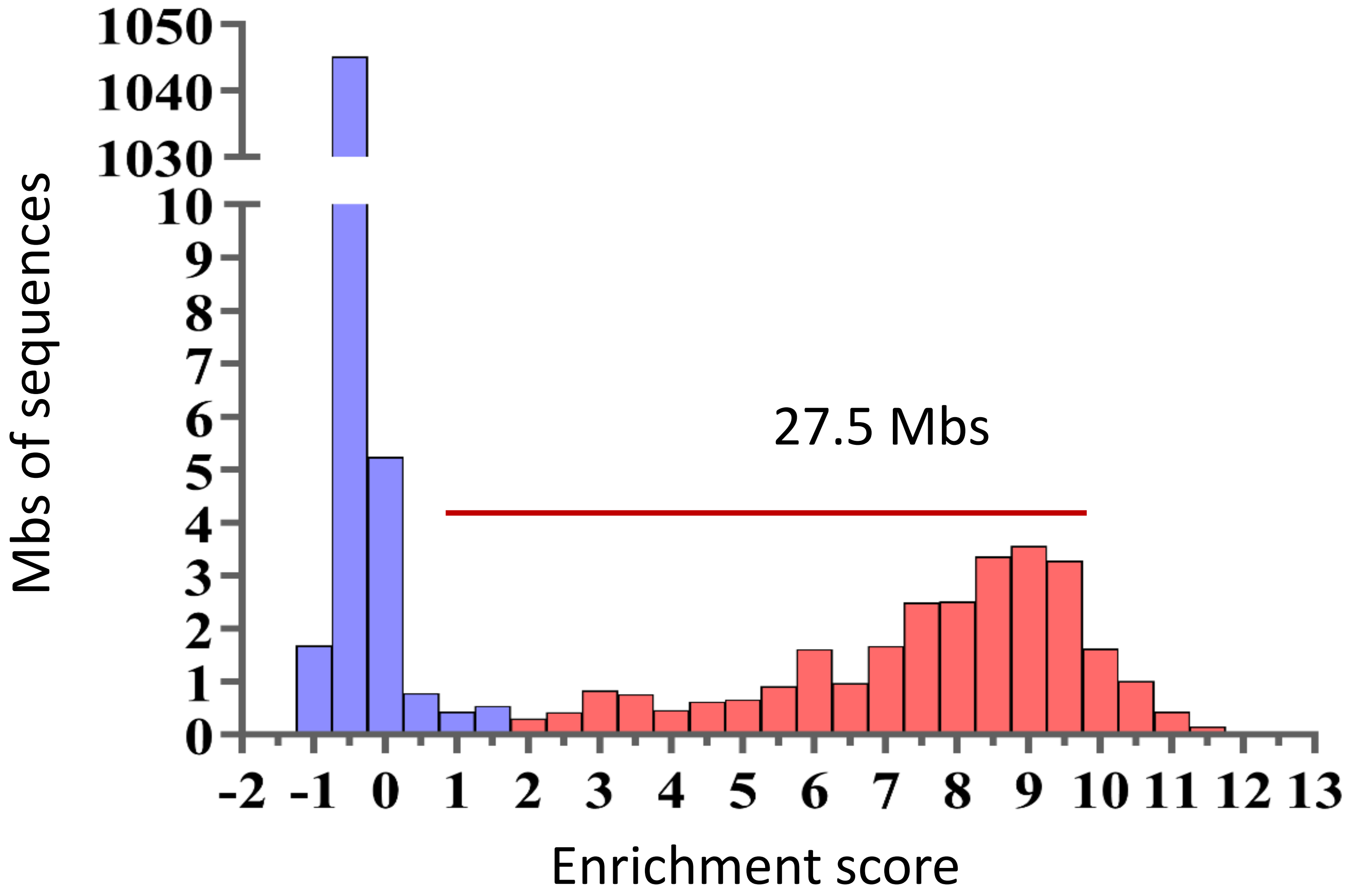


**Supplementary Fig 2:** Distribution of the log2(standardized sperm/blood read coverage): Bars highlighted in red correspond to regions (in Mbps) with 2-fold or greater read coverage in sperm compared to blood (assigned to GSR); blue bars to regions putatively in the somatic genome (coverage <2).

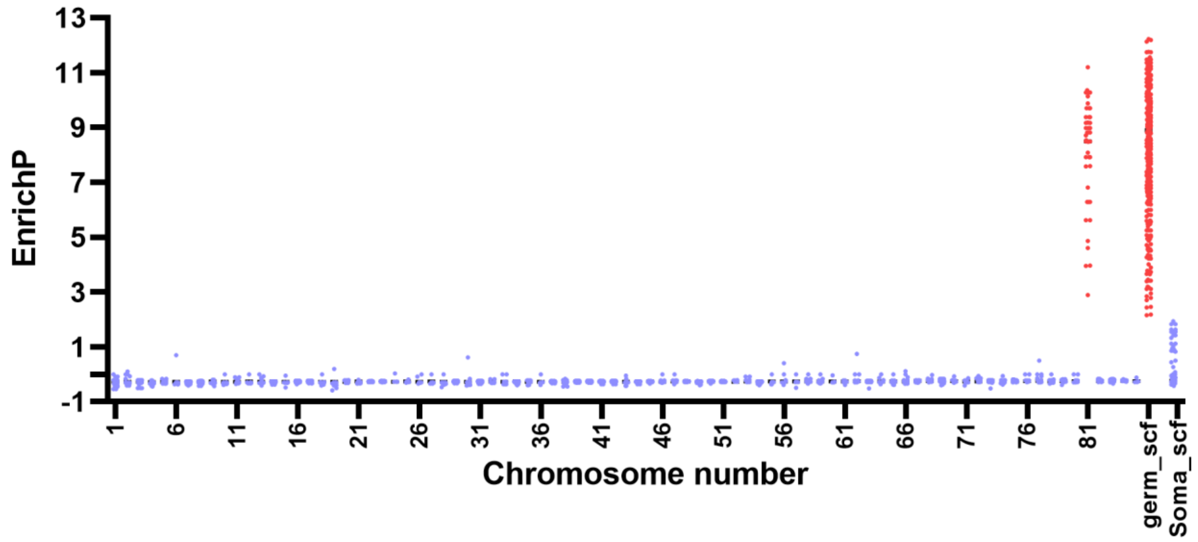


**Supplementary Fig. 3:** Distribution of enrichment score across genome. X-axis represents the chromosome locations and Y axis represents the the enrichment score in each chromosomal region. Chromosome 81 and germ_scf, highlighted in red, represent the region where the enrichment score was 2 or higher. If the enrichment score is less than 2, it is highlighted in purple.

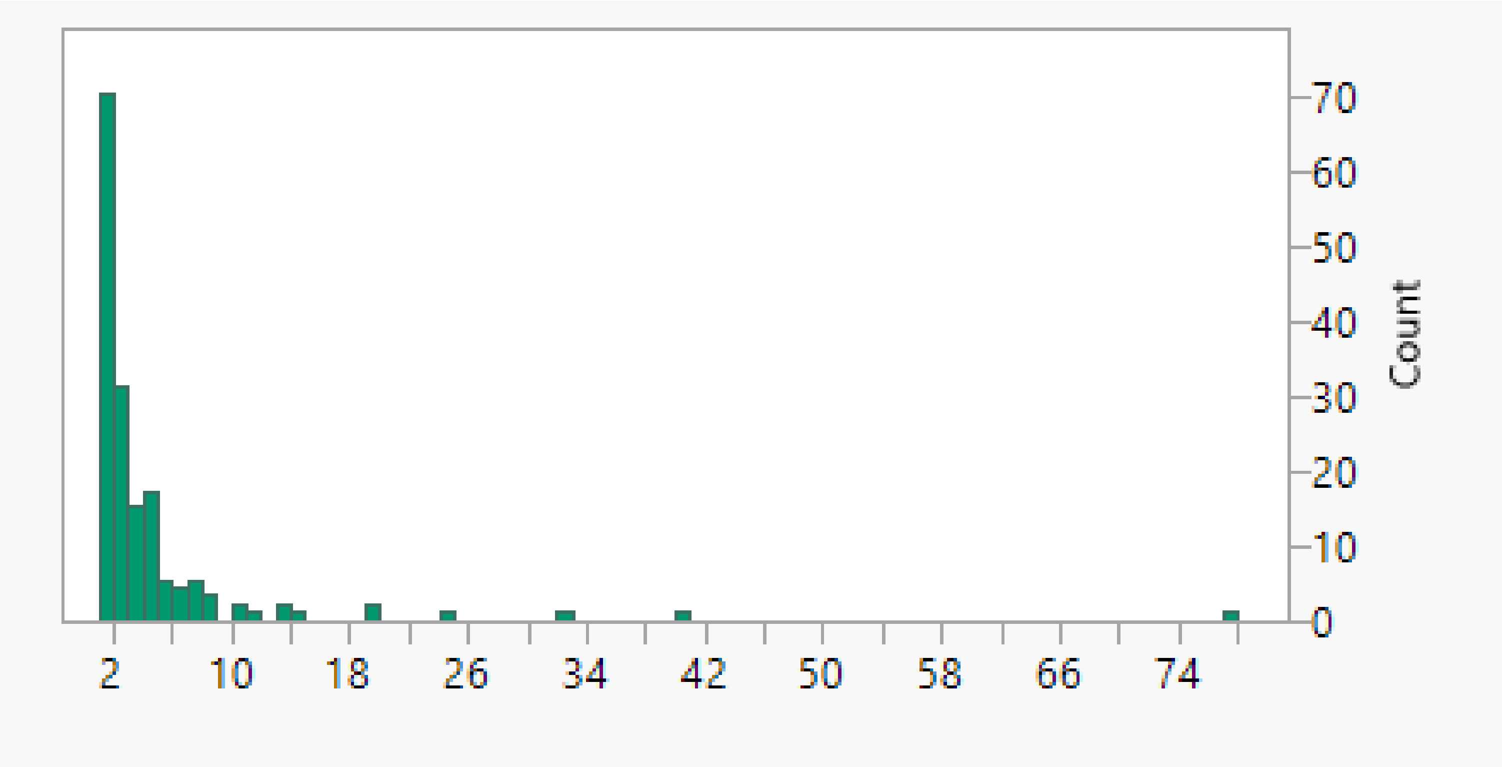


**Supplementary Fig. 4: Histogram of the number of copies of GSGs in the GSR.** X-axis is the frequency of a given GSGs in GSR while the Y-axis is the number of GSGs with that frequency.

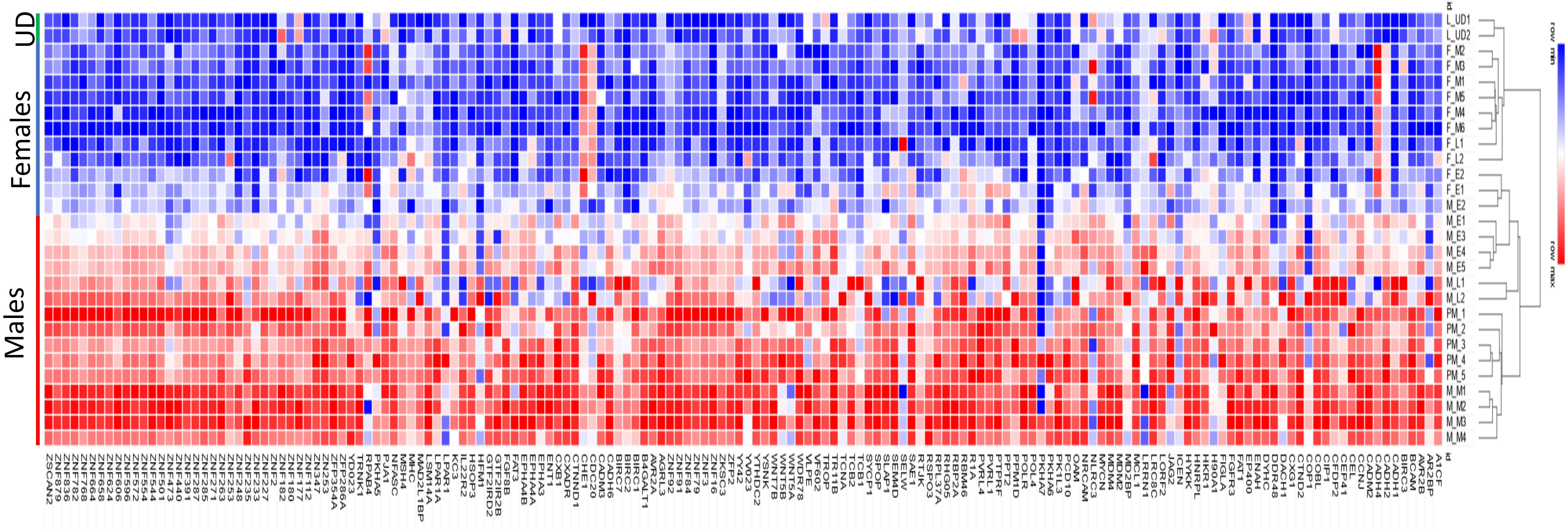


**Supplementary Fig. 5:** Heatmap for all GSGs in 28 gonad samples showing differences in expression among stages. UD stands for undifferentiated larvae

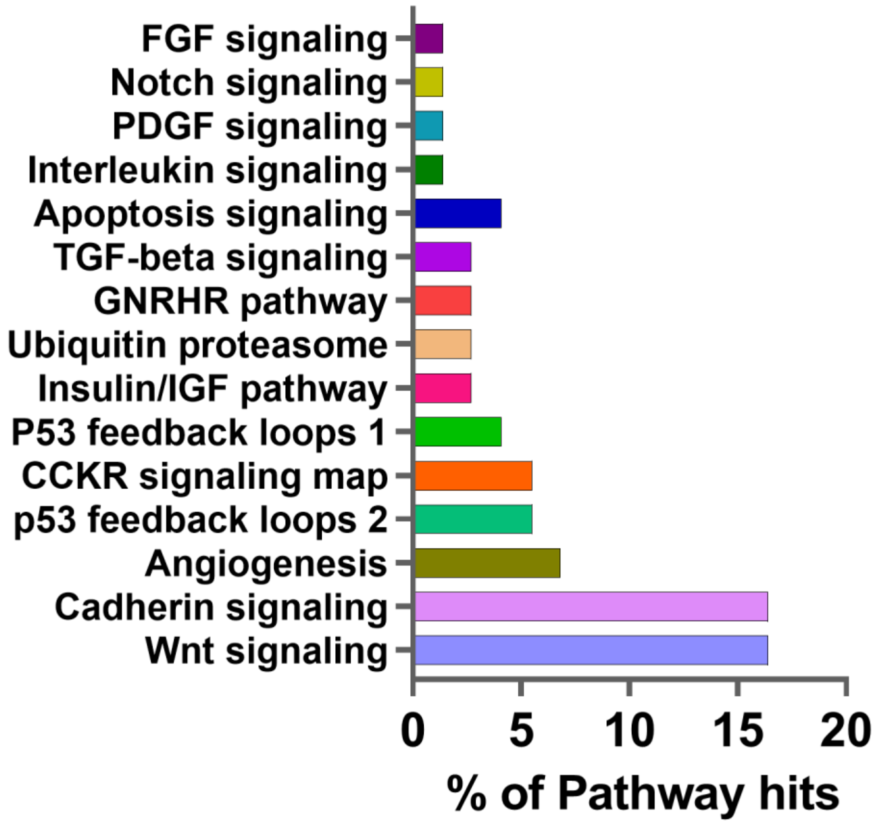


**Supplementary Fig. 6:** A histogram showing the pathways in which GSGs are involved along with the percentage of pathway hits. X-axis represents the percentage (%) of pathway hits and Y-axis represents different pathways involved

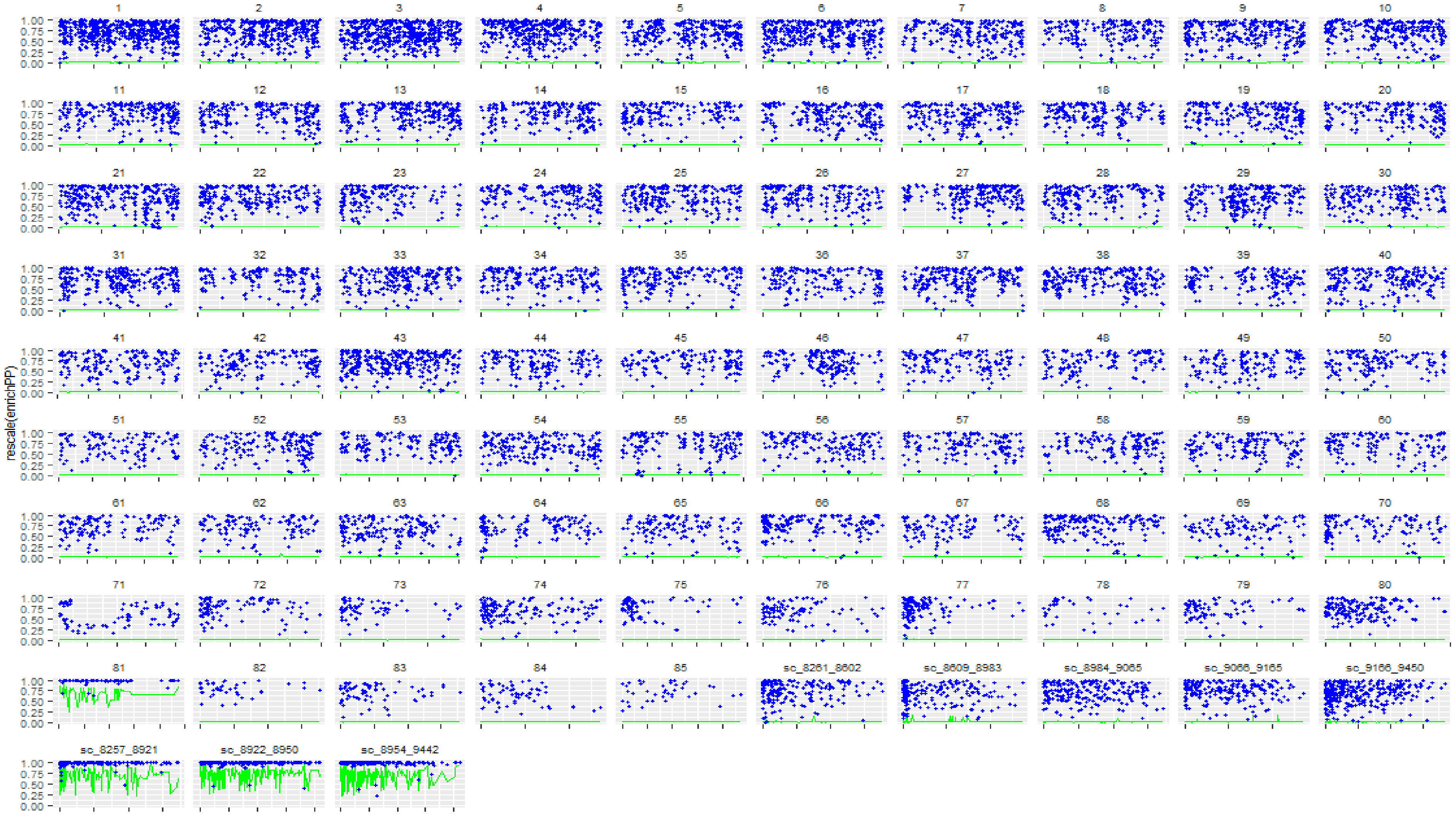


**Supplementary Fig. 7:** Proportion of male-biased genes in different chromosomal regions of sea lamprey genome. Y axis represents the proportion in male vs female. The proportion scale 0-1 refers to the scale of male-biasness compared to females. X axis represents the genes present in those regions. The green line represents the enrichment score of each gene across genome.

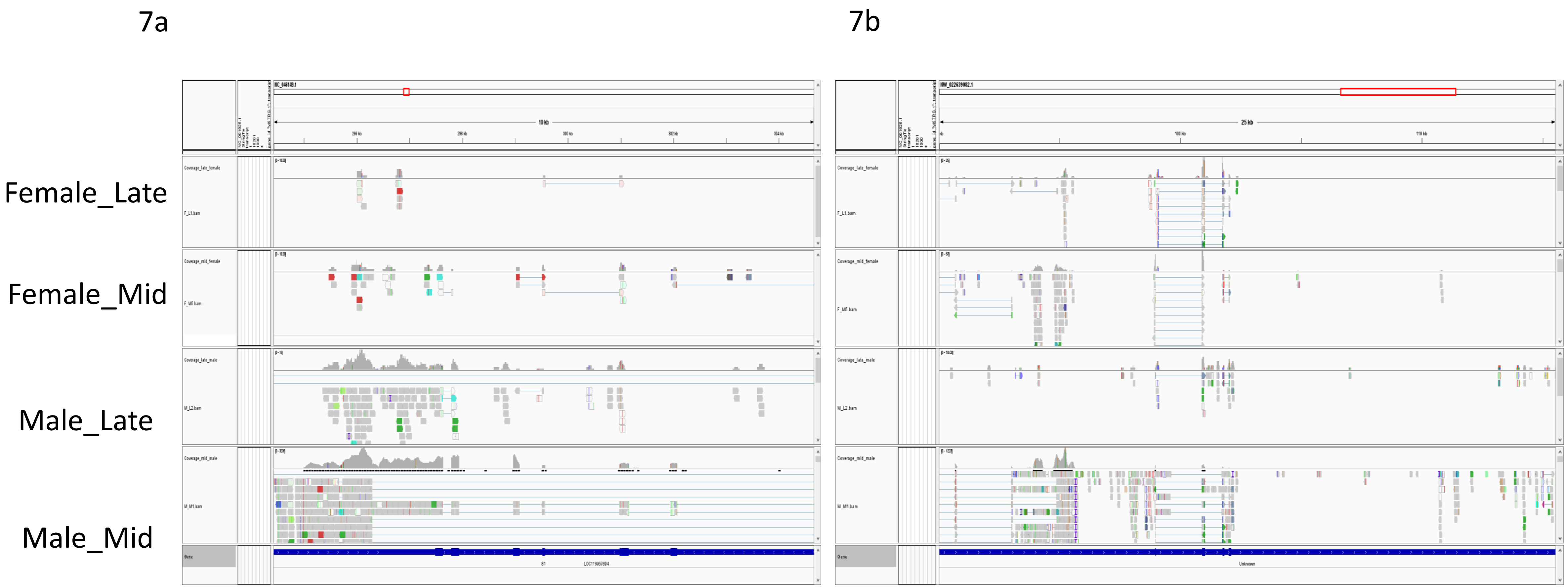


**Supplementary Fig. 8:** Germline-specific genes and their expression in late and mid female and males present in germline-specific region in chromosome 81 and in unplaced scaffolds in the genome viewed in IGV.
a) gene LOC116957694 b) gene LOC116937865

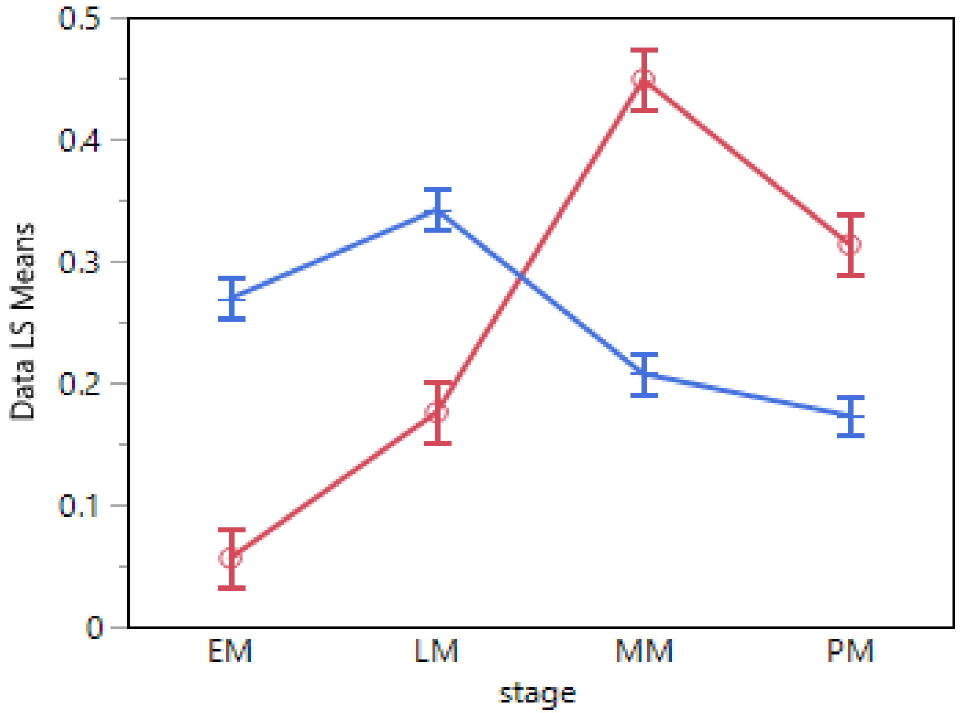


**Supplementary Fig. 9:** Least squares means plot showing differences in expression of GSG paralogues across genome and life stages. X axis represents different stages of male gonadal development where EM denotes early males, LM for late males, MM for mid males and PM for prospective males. Y axis represents Least square means.

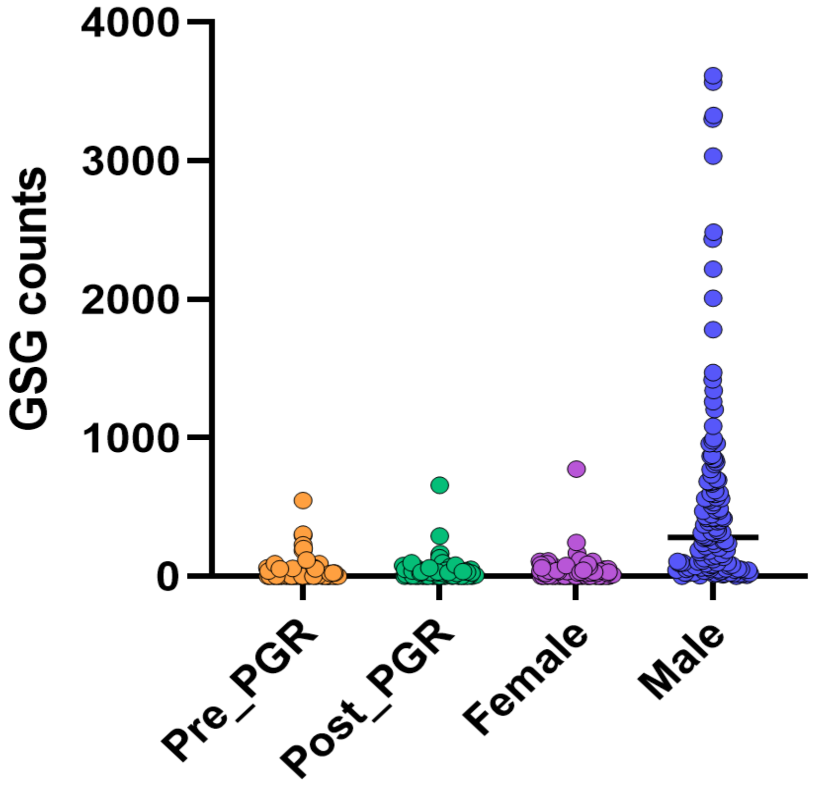


**Supplementary Fig. 10:** Expression of GSGs present in pre-PGR and post-PGR embryo and their comparison across stage and sexes. Y axis presents number of GSG counts and X axis presents pre- and post-PGR stages, males and females. Pre-PGRs are 1, 2, 2.5 dpf embryos and Post-PGR are 3, 4, 5 dpf embryos

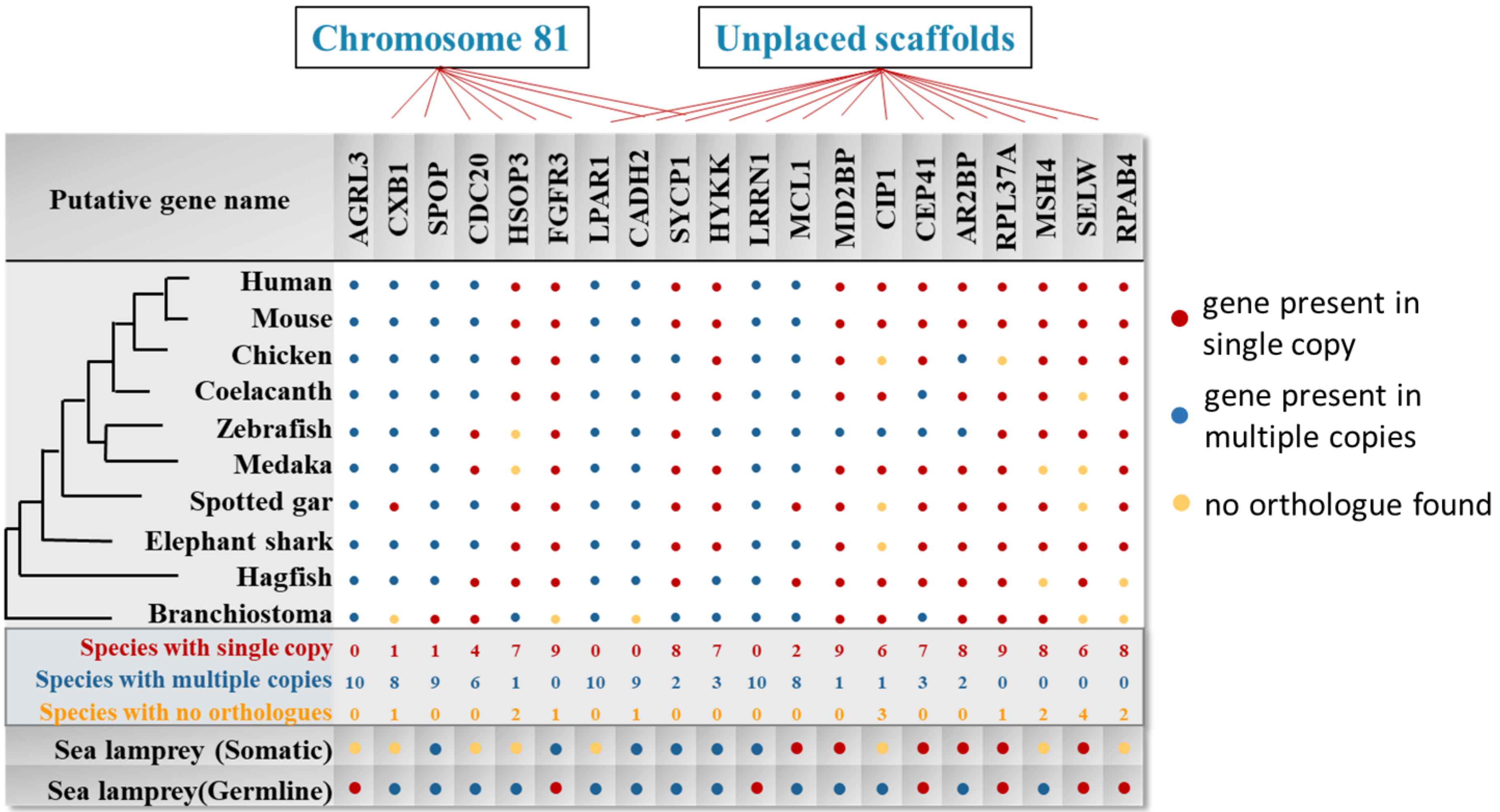


**Supplementary Fig 11**. Evolutionary conservation of copy number of GSGs in the 11 species; putative gene names are given across the top, with the genomic location of the genes in the sea lamprey germline genome and gene names below.

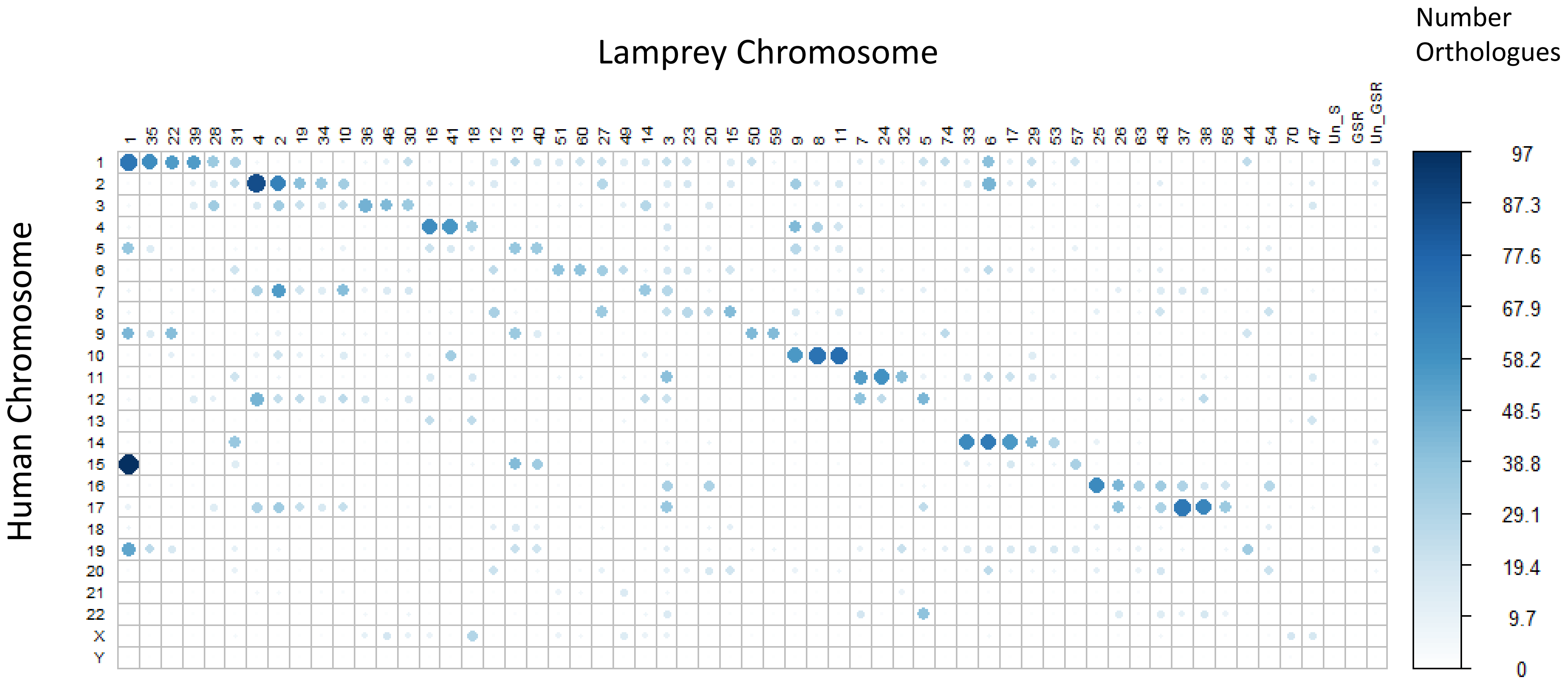


**Supplementary Fig. 12:** Synteny plot showing comparative mapping in sea lamprey and human genome

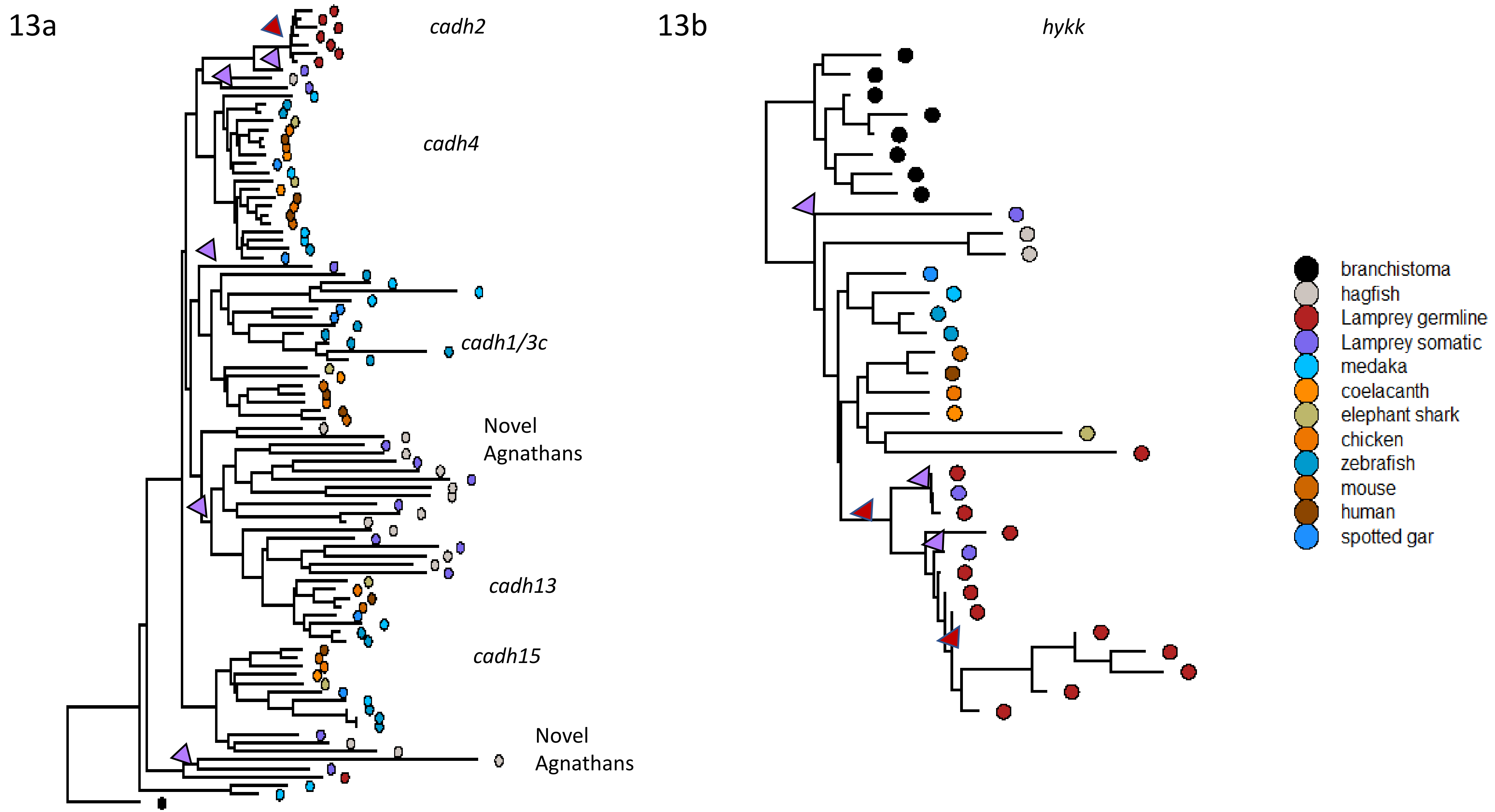
13a
cadh2
cadh4
cadh1/3c
Novel Agnathans
cadh13
cadh15
Novel Agnathans
13b
hykk
branchistoma
hagfish
Lamprey germline
Lamprey somatic
medaka
coelacanth
elephant shark
chicken
zebrafish
mouse
human
spotted gar

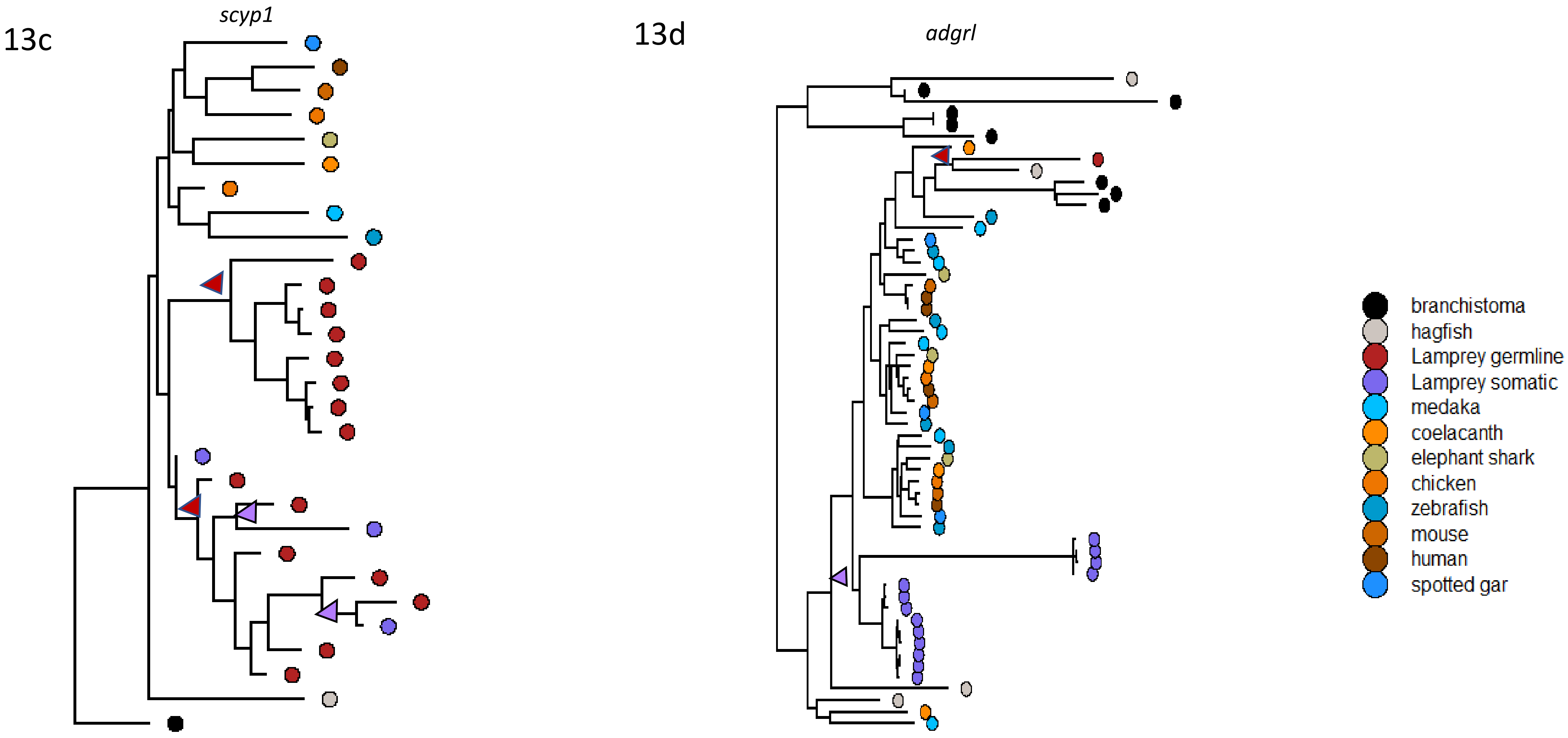


**Supplementary Fig. 13:** Phylogenetic tree for the gene a) *cadh*, b) *hykk*, c) *sycp1*, and d) *adgrl*. Green triangle indicates the place where a somatic paralogue of the gene is present and red triangle indicates the place where germline paralogue of the gene is present.

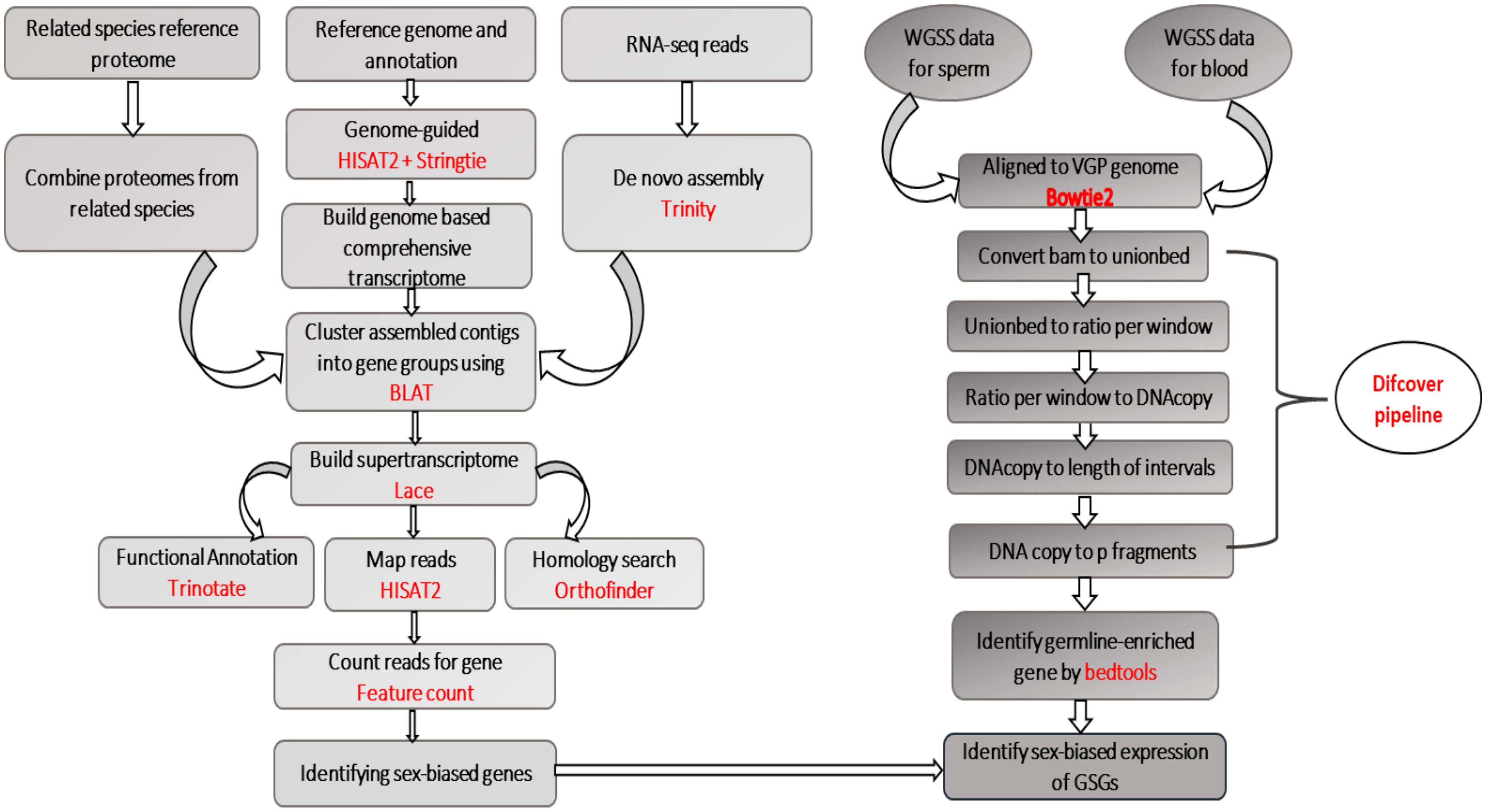


**Supplementary Fig. 14:** Step-by-step workflow and pipelines used for the study.

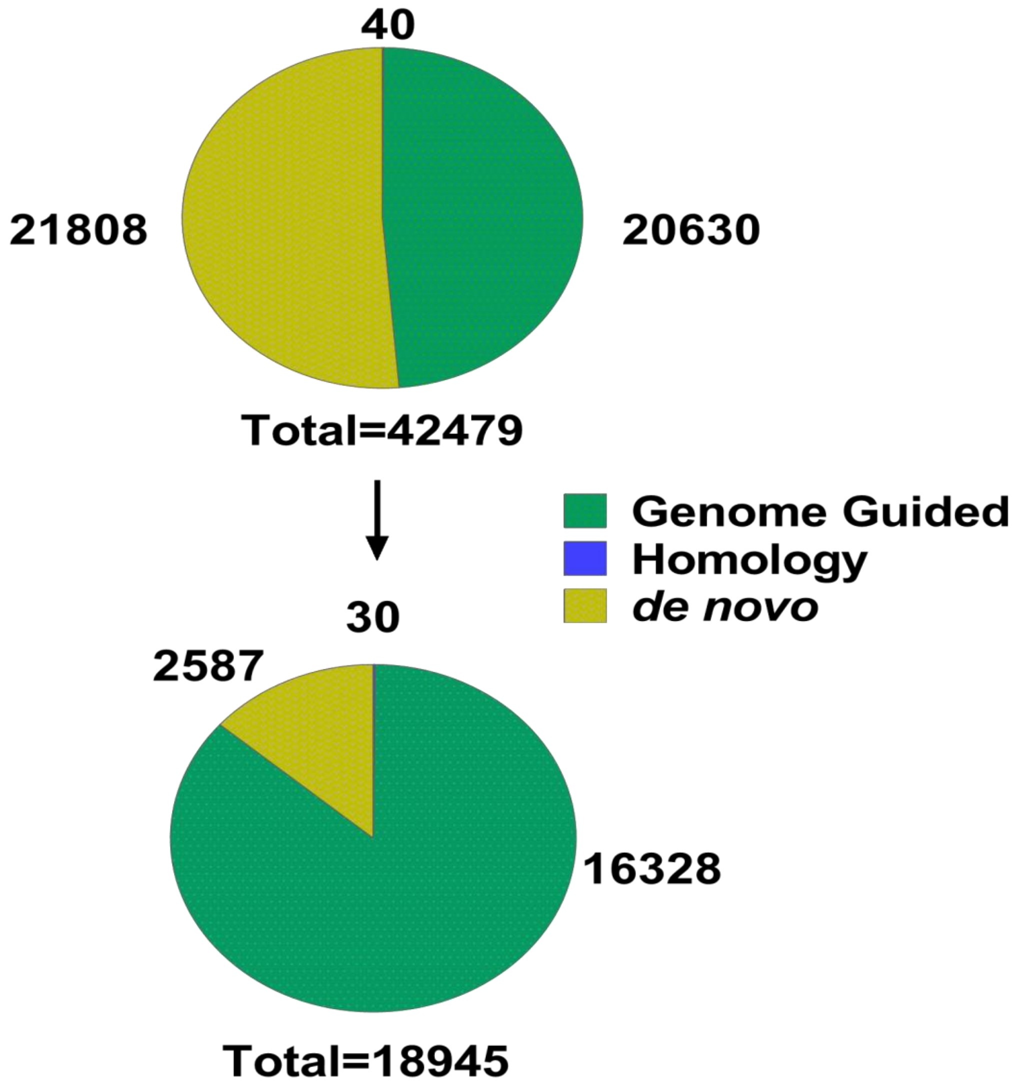


**Supplementary Fig. 15:** Pie charts showing the number of genes identified by the three-tiered necklace pipeline. Top pie chart showing the initial number of genes generated by the pipeline and bottom chart showing the final number of post-filtering genes used for the study.

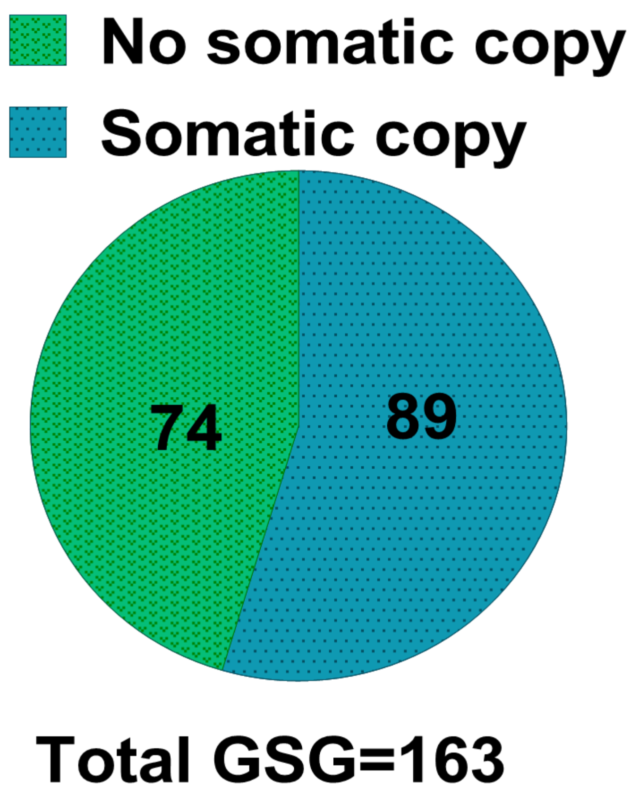


**Supplementary Fig. 16:** Pie charts showing the number of somatic and germline paralogues of GSGs

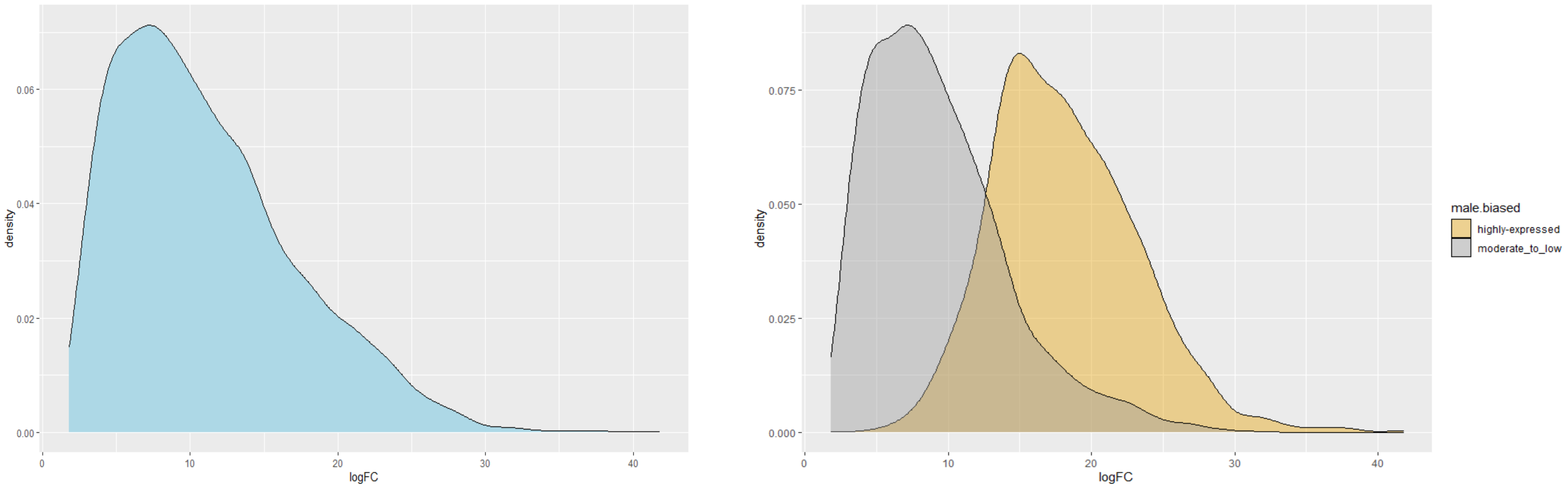


**Supplementary Fig. 17:** Density plots showing the overall logFC (log fold change) a) all male-biased genes in the genome b) differences in density according to the level of biasness in expression in males
