## Supplementary Table 1 & 13 for "The germline-specific region of the sea lamprey genome plays a key role in spermatogenesis"

**Supplementary Table 1:** Details of sea lamprey (*Petromyzon marinus*) samples used in this study. Sex was determined by visual inspection of the gonad during dissection, stages of metamorphosis were identified according to the morphological criteria outlined in[1], and gonadal characteristics were inferred from sex, larval size, and life stage (see Supplementary Fig. 1[2]).

| Sex | Stage | Sample ID | Sample Date | Length (mm) | Collection Site | Basin | Life Stage | Gonadal Characteristics |
|---|---|---|---|---|---|---|---|---|
| **Undetermined** | **Undifferentiated Larvae** | L_UD1 | July 2016 | 57 | Au Sable R, MI | Huron | Larval stage | Small, histologically undifferentiated gonad |
| | | L_UD2 | April 2018 | 66 | Richibucto R, NB | Atlantic | Larval stage | “ ” |
| **Female** | **Early Female** | F_E1 | April 2018 | 81 | Chippewa R, MI | Huron | Larval stage | Ovarian differentiation in progress/completed; synchronous production of primary oocytes with the onset of meiosis I; majority of germ cells have become oocytes |
| | | F_E2 | April 2018 | 95 | Chippewa R, MI | Huron | Larval stage | “ ” |
| | **Mid Female** | F_M1 | Aug 2015 | 135 | Richibucto R, NB | Atlantic | Metamorphosing stage 1 | Oocytes arrested in meiotic prophase; cytoplasm grows at gradual rate |
| | | F_M2 | Aug 2015 | 130 | Richibucto R, NB | Atlantic | Metamorphosing stage 1 | “ ” |
| | | F_M3 | July 2016 | 128 | Richibucto R, NB | Atlantic | Metamorphosing stage 3 | “ ” |
| | | F_M4 | Aug 2017 | 120 | Richibucto R, NB | Atlantic | Metamorphosing stage 4 | “ ” |
| | | F_M5 | Nov 2015 | 129 | | Huron or Michigan[1] | Metamorphosing stage 7 | “ ” |
| | | F_M6 | Oct 2017 | 130 | Richibucto R, NB | Atlantic | Post-metamorphosis | “ ” |
| | **Late Female** | F_L1 | June 2018 | 510 | Ocqueoc R, MI | Huron | Late upstream migrant | Late sexual maturation; vitellogenesis complete, ovulation approaching/complete |
| | | F_L2 | June 2018 | 550 | Ocqueoc R, MI | Huron | Late upstream migrant | “ ” |
| **Prospective Male** | **Prospective Male** | PM_1 | April 2018 | 74 | Richibucto R, NB | Atlantic | Larval stage | Small, histologically undifferentiated gonad beyond the size at which ovarian differentiation is in progress/complete |
| | | PM_2 | April 2018 | 75 | Chippewa R, MI | Huron | Larval stage | “ ” |
| | | PM_3 | July 2018 | 82 | Chippewa R, MI | Huron | Larval stage | “ ” |
| | | PM_4 | July 2018 | 99 | Chippewa R, MI | Huron | Larval stage | “ ” |
| | | PM_4 | July 2017 | 118 | Richibucto R, NB | Atlantic | Larval stage/ Metamorphosing stage 1 | “ ” |
| **Male** | **Early Male** | M_E1 | July 2017 | 132 | Richibucto R, NB | Atlantic | Metamorphosing stage 1 | Early stage of spermatogonial differentiation and production of Type A spermatogonia |
| | | M_E2 | Aug 2017 | 129 | Richibucto R, NB | Atlantic | Metamorphosing stage 3 | “ ” |
| | | M_E3 | Aug 2017 | 114 | Richibucto R, NB | Atlantic | Metamorphosing stage 4 | “ ” |
| | | M_E4 | Aug 2017 | 126 | Richibucto R, NB | Atlantic | Metamorphosing stage 5 | “ ” |
| | | M_E5 | Aug 2017 | 123 | Richibucto R, NB | Atlantic | Metamorphosing stage 6 | “ ” |
| | **Mid Male** | M_M1 | Nov 2015 | 122 | | Huron or Michigan[1] | Metamorphosing stage 7 | Undergoing spermatogonial proliferation and production of Type A and Type B spermatogonia |
| | | M_M2 | Nov 2015 | 133 | | Huron or Michigan[1] | Metamorphosing stage 7 | “ ” |
| | | M_M3 | Dec 2017 | 130 | Richibucto R, NB | Atlantic | Post-metamorphosis | “ ” |
| | | M_M4 | Dec 2016 | 114 | Richibucto R, NB | Atlantic | Post-metamorphosis | “ ” |
| | **Late Male** | M_L1 | April 2018 | 385 | Black Mallard R, MI | Huron | Early upstream migrant | Early sexual maturation; spermatids, immature sperm |
| | | M_L2 | June 2018 | 400 | Ocqueoc R, MI | Huron | Late upstream migrant | Late sexual maturation; mature sperm, spermiation approaching/complete |

[1] Larval lampreys collected from multiple tributaries of Lake Huron and Lake Michigan and housed communally so specific sample site is not available.

**Supplementary Table 13:** List of sea lamprey GSGs and their tissue of bias that are associated with gonadal development, differentiation or sex determination in other taxa. NB denotes a lack of bias towards male or female.

| Putative gene name | Bias: testes/ovary | Known function in gonad | Species | Reference |
|---|---|---|---|---|
| *AGRL3* | Testes | Female-biased expression | Tilapia | [3] |
| *AR2BP* | NB | | | |
| *AVR2B* | Testes | | | |
| *CADH* | Testes | | | |
| *CADM2/3* | Testes | | | |
| *CFDP2* | NB | | | |
| *EPHA3* | Testes | | | |
| *HNRPL* | Testes | | | |
| *JAG2* | NB | | | |
| *LORF2* | Testes | | | |
| *LRRN1* | NB | | | |
| *NLRC3* | NB | | | |
| *PCD10* | Testes | | | |
| *PKHA5/6/7* | NB | | | |
| *RBM46* | Testes | | | |
| *SEM4D* | Testes | | | |
| *PPM1D* | Testes | Male-biased expression | Tilapia | [3] |
| *CCNJ* | Testes | | | |
| *WNT5A/5B* | Testes | | | |
| *BIRC1* | Testes | Gonad development | Tilapia | [4] |
| *CDC20* | NB | Gonad development | Carp | [4] |
| *AVR2A* | Testes | | | |
| *DDR48* | NB | | | |
| *RBP2A* | Testes | | | |
| *WDR78* | Testes | | | |
| *MYCN* | Testes | | | |
| *GTF2IRD2* | Testes | | | |
| *B4GALT1* | Testes | Female-biased expression | Carp | [3] |
| *DACH1* | NB | Gonad development | Drosophila | |
| *FAT1/3* | Testes | Female-biased expression | Mouse | [5] |
| *MSH4* | Testes | Gonad development | Mouse | [5] |
| *SYCP1* | Testes | | | |
| *FGF8B* | Testes | Sex determination & differentiation | Mouse | [6–8] |
| *RSPO3* | Testes | Sex differentiation | Mammals | [6–8] |
| *FGFR3* | Testes | Sex determination and differentiation | Sturgeon | [9] |
| *H90A1* | NB | Sex-biased expression in male | Olive flounder | [10] |
| *PTPRF* | Testes | Gonad development | Medaka | [11] |
| *PVRL1/4* | NB | | | |
| *RPAB4* | NB | Gonad differentiation | Yellow carp | [12] |
